# BANCOVID, the first D614G variant mRNA-based vaccine candidate against SARS-CoV-2 elicits neutralizing antibody and balanced cellular immune response

**DOI:** 10.1101/2020.09.29.319061

**Authors:** Juwel Chandra Baray, Md. Maksudur Rahman Khan, Asif Mahmud, Md. Jikrul Islam, Sanat Myti, Md. Rostum Ali, Md. Enamul Haq Sarker, Samir Kumar, Md. Mobarak Hossain Chowdhury, Rony Roy, Faqrul Islam, Uttam Barman, Habiba Khan, Sourav Chakraborty, Md. Manik Hossain, Md. Mashfiqur Rahman Chowdhury, Polash Ghosh, Mohammad Mohiuddin, Naznin Sultana, Kakon Nag

## Abstract

Effective vaccine against SARS-CoV-2 is the utmost importance in the current world. More than 1 million deaths are accounted for relevant pandemic disease COVID-19. Recent data showed that D614G genotype of the virus is highly infectious and responsible for almost all infection for 2^nd^ wave. Despite of multiple vaccine development initiatives, there are currently no report that has addressed this critical variant D614G as vaccine candidate. Here we report the development of an mRNA-LNP vaccine considering the D614G variant and characterization of the vaccine in preclinical trial. The surface plasmon resonance (SPR) data with spike protein as probe and competitive neutralization with RBD and S2 domain revealed that immunization generated specific antibody pools against the whole extracellular domain (RBD and S2) of the spike protein. The anti-sera and purified IgGs from immunized mice on day 7 and 14 neutralized SARS-CoV-2 pseudovirus in ACE2-expressing HEK293 cells in a dose dependent manner. Importantly, immunization protected mice lungs from pseudovirus entry and cytopathy. The immunologic responses have been implicated by a balanced and stable population of CD4^+^ cells with a Th1 bias. The IgG2a to IgG1 and (IgG2a+IgG2b) to (IgG1+IgG3) ratios were found 1±0.2 and 1.24±0.1, respectively. These values are comparatively higher than relevant values for other published SARS-CoV-2 vaccine in development,^1, 2^ and suggesting higher viral clearance capacity for our vaccine. The data suggested great promise for immediate translation of the technology to the clinic.

## Introduction

A new infectious corona virus (SARS-CoV-2) has been first reported from Wuhan, China in December, 2019 that causes COVID 19.^3^ The World Health Organization (WHO) declared the COVID-19 a global public health emergency situation on February 5, 2020 after getting growing evidence of continuous person-to-person transmission.^4^ The virus has been spread worldwide quickly, and consequently WHO has declared it pandemic in March 11, 2020. As of September 29, 2020, the pandemic has resulted in 1,007,887,415 deaths among over 33,630,004 patients in 215 countries, with a case-fatality rate of 3%.

There will be a risk of pandemic as long as there is COVID-19 epidemic situation in any area of the world unless people are properly vaccinated. Therefore, effective vaccines against SARS-CoV-2 are immediately required to control morbidity and mortality related with COVID-19. Generally, non-replicating viral vectors, inactivated virus, DNA-based and protein-based vaccines have been the major approaches for the development of stable and effective vaccines; though they have their inherent limitations.^5^ Recently, mRNA-based vaccines have become a promising approach because of their opportunity for rapid development, comparative low dose, logical better safety profile, and low capital expenditure (CAPEX).^6^ Several other leading vaccines under development against SARS-CoV-2 are also mRNA-based.^2,7,8,9^ Lipid nanoparticle technology has been developed for effective delivery of single-stranded therapeutics like siRNA, antisense oligo, mRNA etc. The first RNA-LNP therapeutic was approved in 2018 and has set the example for clinical safety of LNP-formulated RNA.^10^ Therefore, we have also opted for mRNA-based LNP-mediated vaccine development technology to support the initiative for preventing the ongoing wave of the COVID-19 pandemic.

The candidate mRNA vaccine ‘BANCOVID’ is a LNP–encapsulated, nucleoside-modified mRNA–based vaccine that encodes the SARS-CoV-2 spike (S) glycoprotein stabilized in its prefusion conformation. Coronaviruses have genetic proofreading mechanisms, and SARS-CoV-2 sequence diversity is comparatively low;^11, 12^ though, natural selection can adopt rare but favorable mutations. Since the outbreak in China, SARS-CoV-2 has gone through numerous mutations. Among these, the D614G amino acid change in the spike protein of Wuhan reference strain is caused by an A-to-G nucleotide substitution at position 23,403 of the relevant nucleotide sequence. Currently, D614G is the most prevalent circulating isotype of SARS-CoV-2 worldwide (more than 95%).^13, 14^ To date, there is no published report about the D614G-relevant vaccine development. Few studies have shown that antibody generated using D614 variant-target did not show significant difference between D614 and G614 variants in terms of cellular entry.^15^ These studies did not use G614-specific antibody, and applied artificial systems for characterizing relevant functional experiments. Furthermore, how G614 variant vaccine behave in immunization and what would be the impact of relevant antibody on SARS-CoV-2 is not known. Therefore, developing of G614 variant-specific vaccine is a prime importance, and warrant characterization. To address this, we have incorporated D614G variant-targeted nucleic acid sequence, as well as few other immunogen-enhancing aspects in our mRNA design consideration. In this study, we described the design and preclinical characterization of ‘BANCOVID’ mRNA-LNP vaccine candidate.

## Materials and Methods

### Target gene and vector cloning

#### Target selection

As of March 27, 2020, there were 170 surface glycoproteins (partial and complete sequence) out of 1661 SARS-CoV-2 proteins posted on NCBI Virus database. A comparative sequence alignment using Clustal Omega (https://www.ebi.ac.uk/Tools/msa/clustalo/) showed differences in several regions, notably in position 614 (D>G). A total of 15 glycine containing surface glycoprotein were found instead of aspartic acid. A consensus sequence from multiple sequence alignment was identified (data not shown) using EMBOSS Cons (https://www.ebi.ac.uk/Tools/msa/emboss_cons/) and selected as primary target sequence for vaccine development. Hydrophilicity/hydrophobicity plot analysis was performed using GENETYX Ver8.2.0, protein 3D modeling using Phyre2^16^ and visualized using UCSF Chimera 1.11.2rc.^17^ Finally, D614G mutation and double proline (2P) mutations (K986P and V987P) were incorporated to the target sequence.

#### Target amplification

Nasopharyngeal and oropharyngeal swab sample were collected from a COVID-19 positive male patient. Virus heat inactivation at 56 °C for 30 minutes and total RNA including virus RNA extraction was performed using TRIzol™ Plus RNA Purification Kit (ThermoFisher, USA). cDNA synthesis was performed using GoScript™ Reverse Transcription System (Promega, USA). S-gene (Surface glycoprotein) was amplified using 3 different sets of primers, 0572F and 0573R, 0574F and 0575R, 0576F and 0577R, respectively (supplementary table 1) and Platinum™ SuperFi™ DNA Polymerase (ThermoFisher, USA). Amplified S-gene and polymerase chain reaction (PCR) engineered pET31b(+) (Novagen, Germany) bacterial expression vector were amplified using 0570F and 0571R primers, excised and extracted from agarose gel using GeneJET Gel Extraction and DNA Cleanup Micro Kit (ThermoFisher, USA), and assembled together using NEBuilder® HiFi DNA Assembly Master Mix (NEB, USA). Sub-cloning was performed into DH5alpha chemical competent cells, miniprep purification was using PureLink™ Quick Plasmid Miniprep Kit (ThermoFisher, USA). S-gene integration check into vector was performed via restriction digestion using XbaI (ThermoFisher, USA) and EcoRI (ThermoFisher, USA), and PCR using primers 0600F and 0024R (supplementary table. 1). DNA sequencing was performed to confirm the complete open reading frame (ORF) compatibility of target S-gene. Finally, sequence confirmed rDNA (rDNA ID: p20004, supplementary figure 2) was further amplified and purified using PureLink™ HiPure Plasmid Midiprep Kit (ThermoFisher, USA), sequenced, and stored for future purposes. Also, sequence confirmed S-gene was submitted to NCBI (GenBank accession number MT676411.1), where we identified and noted D614G mutation. Supplier’s manual with minor modifications were followed for all the methods.

**Table 1:**
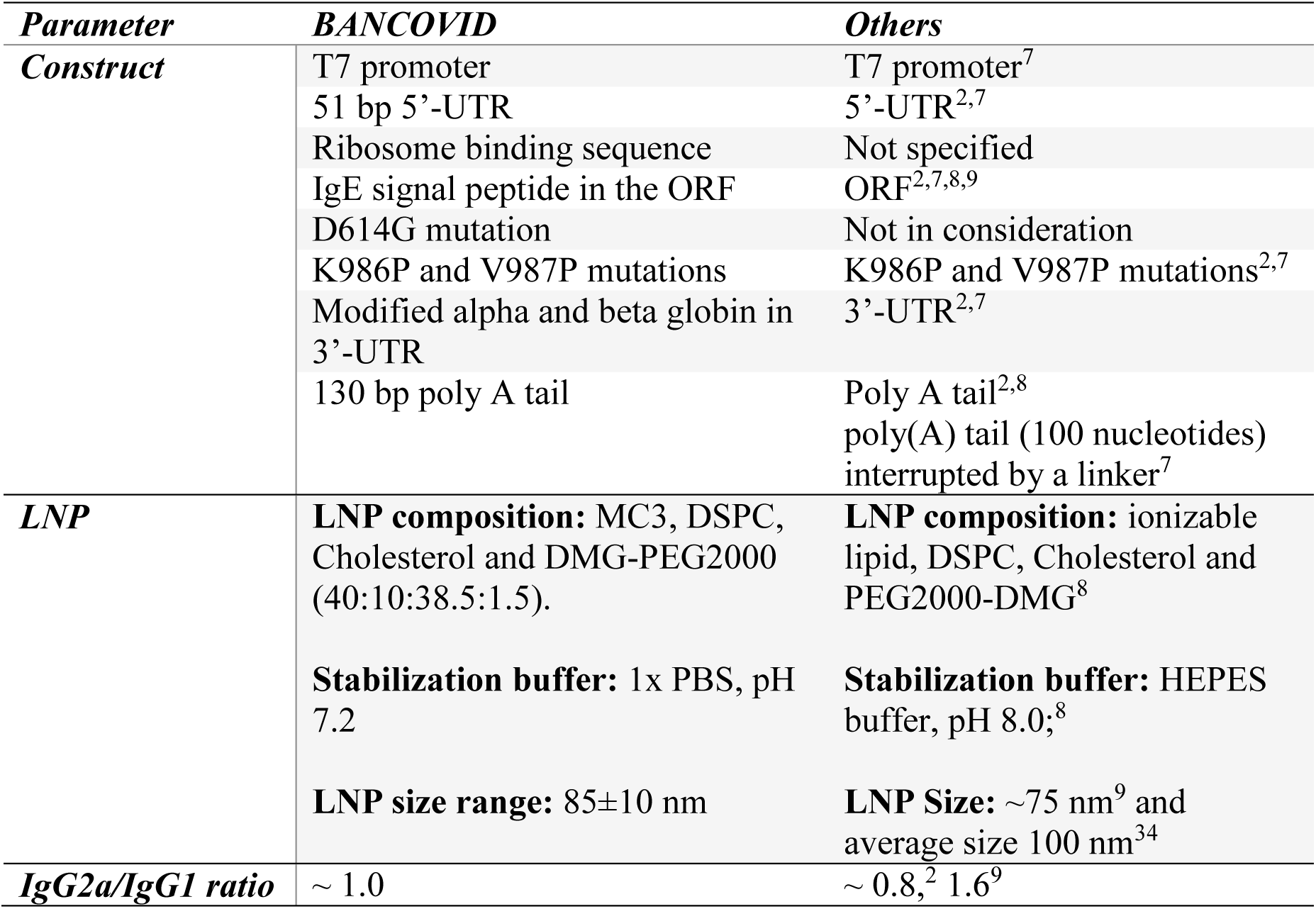
Comparative design features of ‘BANACOVID’.

#### Target modification

An immunoglobulin (Ig) heavy chain (HC) 19 amino acid signal peptide (H1)^18^ was assembled (0583F, 0584R, 0585F and 0586R), and amplified (0583F and 0586R) along with homology arm for incorporating into rDNA p20004, replacing native 13 amino acid leader sequence. Assembled signal peptide was amplified with homology arm and rDNA p20004 was engineered via PCR using 0582F and 0571R primers (supplementary table 1 for assembly and amplification primers). New rDNA p20006 (data not shown) was prepared by incorporating signal peptide and engineered p20004 rDNA, using above explained method as p20004 rDNA preparation.

S-gene was amplified from rDNA p20006 using 0594F and 0592RR primers (supplementary table 1). This gene and pcDNA™5/FRT Mammalian Expression Vector (ThermoFisher, USA) were digested using Acc65I (ThermoFisher, USA) and XhoI (ThermoFisher, USA) and visualized via agarose gel electrophoresis. The desired bands from the gel were excised and purified using GeneJET Gel Extraction and DNA Cleanup Micro Kit and ligated using T4 DNA Ligase (ThermoFisher, USA). After ligation, sub-cloning into DH5α chemical competent cells, plasmid miniprep purification, insert checking, DNA sequencing, plasmid midiprep purification, DNA sequencing and storage (rDNA ID: p20010, data not shown) were performed.

2P (double Proline) amino acid mutations at position 986 (K986P) and 987 (V987P) were also performed via site directed mutagenesis using 0745F and 0745R primers (supplementary table 1). DNA sequencing was performed to confirm desired mutations (rDNA ID: p20015, data not shown).

Finally, a T7 promoter sequence, a synthetic 5’-UTR, an IgE signal peptide replacing native 13 amino acids signal peptide from S-gene, a 3’-UTR (modified alpha globin and modified beta globin), and a 130 bp synthetic poly A-tail (pA-tail) were added. A restriction endonuclease (Sfo I) sequence before T7 promoter sequence and after pA were added for cutting out desired size of DNA for in-vitro mRNA synthesis. Final rDNA ID was p20020 (supplementary figure 2) and rDNA construction was performed as mentioned before, same as p20004 and p20006 (supplementary figure 2). Supplier’s manual with minor modifications were followed for all the methods.

#### Sequencing

DNA sequencing was performed as according to supplier’s protocol for the final construct p20020 and other constructs e.g., p20004, p20006, p20010, p20015 etc. (supplementary table 2 for sequencing primers) using 3500 Genetic Analyzer (ThermoFisher, USA). DNA sequencing data clearly confirmed the presence of the target sequences and modifications.

BigDye® Terminator v1.1 Cycle Sequencing Kits (ThermoFisher, USA) and POP-6 polymer (ThermoFisher, USA) chemistry was used for DNA sequencing reaction.

### mRNA production

#### mRNA synthesis

The in-vitro (IVT) mRNA synthesis reaction was performed using MEGAscript™ T7 Transcription Kit (ThermoFisher, USA), and Ribonucleotide Solution Set (NEB, USA). During development phase, IVT mRNA synthesis reaction was optimized into 4 steps (supplementary method 1). In optimized IVT mRNA reaction, final concentration of ribonucleotides was as follow: ATP and UTP – 13.13 mM, and GTP and CTP – 9.38 mM. The reaction was run for 2 hours at 37 °C. IVT reaction was followed by a DNase treatment at 37 °C for 15 minutes using TURBO DNase from the transcription kit.

#### mRNA capping and purification

mRNA capping reaction was performed with purified IVT mRNA using 3’-O-Me-m7G(5’)ppp(5’)G RNA Cap Structure Analog (NEB, USA). The reaction condition was followed according to supplier’s manual. During development phase, IVT mRNA was purified via protein degradation by phenol:chloroform:isoamyl alcohol, phenol removal by chloroform (twice), and final purification using MEGAclear™ Transcription Clean-Up Kit (ThermoFisher, USA). Capped mRNA purification was performed using the same cleanup kit. Supplier’s instructions were followed during the purification steps. Purified IVT mRNA and capped mRNA were quantified using Multiskan GO spectrophotometer (ThermoFisher, USA).

### Formulation of mRNA

#### mRNA-LNPs formation

Purified mRNAs were first diluted with sodium acetate buffer at desired concentration. The lipid molecules were dissolved in ethanol and mixed well. Lipids (MC3: DSPC: Cholesterol: DMG-PEG2000) were combined in the molar ratio of 50:10:38.5:1.5. ^19,20,21,22^ Then, sodium acetate buffer containing mRNA and lipid sample were mixed at a ratio of 3:1 and passed through the liposome extruder (Genizer, USA) to encapsulate the mRNA. The size distribution was checked after encapsulation of mRNA into nanoparticles. Then the formulations were dialyzed against 50 mM HEPES/sodium acetate buffer and phosphate-buffered saline for 18 hours. The size distribution was again checked after dialysis by Zetasizer Nano ZSP (Malvern, USA). LNP samples were analyzed for size distribution in 1× phosphate buffered saline (PBS) as dispersant. The formulation was concentrated using Ultra centrifugal filters (Merck, Germany), filtered through 0.22 micron filter, and stored at 5±3 °C.^23^ The formulation was passed through the quality control for the particle size, encapsulation efficiency, endotoxin limit and sterility.

### Safety and efficacy in mice

A total number of 50 BALB/c swiss albino mice (male and female) of 6-8 weeks old, were selected randomly and isolated 5 days before immunization. After careful observation and conditioning, 30 mice (15 males and 15 females) were taken to the experiment room for immunization and subsequent safety and efficacy analysis. 9 male mice were also separated for local tolerance testing. The temperature in the experimental animal room was 26 °C (±2 °C) and the relative humidity was 60±5%. The room was HVAC controlled ISO class 7 room with 70% fresh air intake and full exhaust. The mice were individually housed in polypropylene cage with individual water bottle, provided with 5 g of in-house mouse feed daily and kept under 12 hours of day-night cycle. 30 mice were separated into 5 different groups consisting 6 mice (3 males and 3 females) in each group. There were 3 different treatment groups such as Treatment group 1, 2, and 3, 1 placebo group and 1 control group. Each mouse of treatment groups 1, 2, and 3 was immunized with sterile 0.1 µg/50 µL, 1.0 µg/50 µL and 3.0 µg/50 µL of BANCOVID, respectively. Each mouse of the placebo group was injected with the vehicle only and the control group mice were not injected with anything. Intramuscular (IM) injection in the left quadriceps was done for immunization. The flow of the experimental design is shown in supplementary figure 3. The study plan and procedures were approved by the internal ethical review board, which is complied with local ethical regulation. No treatment randomization and blinding methods were used in the study and sample sizes were determined by the resource equation method.

#### Local tolerance

Local tolerance was confirmed by clinical signs, macroscopic and histopathology evaluations of injection sites in animals. 9 male mice were separated for local tolerance study and divided into 3 different groups consisting of 3 male mice in each group. There was 1 treatment group, 1 placebo group and 1 control group for the study. The treatment group was immunized with IM injection with 3.0 µg/50 µL of BANCOVID in the left quadriceps muscle whereas the placebo group was injected with 50 µL of vehicle and the control group with 50 µL of normal saline. Euthanasia and evaluation of lesions was performed in one representative mouse from placebo and control group and 3 from the treatment group at 48 hours post treatment. The inner thigh muscle of injected site of each mouse was excised and placed in 10% neutral buffered formalin until adequately fixed. After trimming, processing and paraffin embedding, the sections are HE stained and observed for erythema and edema under microscope.

#### Immunogenicity

The immunogenicity of BANCOVID was evaluated in BALB/c mice, post administration to the quadriceps muscle. Approximately 200 µL blood was collected from facial vein and centrifuged at 1500 X g for serum isolation (10 minutes at 4 CC). All serums were aliquot, frozen immediately and stored at −80 °C until analysis. The reactivity of the sera from each group of mice immunized with BANCOVID was measured against SARS-CoV-2 S antigen (SinoBiologicals, China). Analysis revealed IgG binding against SARS-CoV-2 S protein antigens in the sera of the immunized mice. The serum IgG binding endpoint titers (EPTs) were measured in mice immunized with BANCOVID. EPTs were observed in the sera of mice at day 7 and day 14 after immunization with a single dose of the vaccine candidate.

#### Toxicity

Pre-immune whole blood (approximately 50 µL) from each mouse was collected for complete blood count (CBC) in 2% EDTA at 3 days before immunization. Similarly, whole blood was also collected after immunization at day 14 for CBC analysis using auto hematology analyzer BK-6190-Vet (Biobase, China). Pre-immune serum of 3 days before and 14 days after immunization were used for chemistry analysis using semi-automatic chemistry analyzer (Biobase, China) such as alanine transferase (ALT), aspartate transaminase (AST) and blood nitrogen urea (BUN).

### Neutralization assay

#### Pseudovirus preparation

Pseudotyped SARS-CoV-2 adeno virus was prepared expressing the SARS-CoV-2 surface glycoprotein gene (S gene) on the virus. S gene of SARS-CoV-2 was cloned into pAADV-B02 vector (Genemedi, China) that also contains a GFP gene downstream of the gene of interest. After construction, SARS-CoV-2 S gene containing plasmid p20017 and adenovirus backbone plasmid pAADV-C01 (Genemedi, China) were co-transfected into HEK293 based adapted viral production cell (ThermoFisher, USA). Viral production cells were seeded in a 6 well TC treated plate (Nest, China) at a concentration of 6 × 10^5^ cell per well and cultured overnight.

Then co-transfection was performed using Lipofectamine 3000 (ThermoFisher, USA) reagent according to manufacturer’s protocol. Next day 1.25% low melting agarose in DMEM media was spread on the well and incubated until plaques were formed. After formation of plaques, multiple plaques were collected in DMEM media and titers were measured for plaque selection. Then selected plaque was added on the fresh pre seeded viral production cell. After few days, cells and supernatant were collected and performed repeated freeze thawing for collection of viruses (P1 pseudovirus). Similarly, infection was performed on fresh cells and virus was collected (P2 pseudovirus). These processes were repeatedly performed and P4 pseudoviruses were collected. After collection of P4 pseudoviruses, concentration and purification was performed by ultracentrifugation and sucrose gradient.^24^ After titer determination, pseudoviruses were stored at −86 °C freezer (ThermoFisher, USA).

Another Pseudotyped SARS-CoV-2 retro virus was prepared that virus have SARS-CoV-2 surface glycoprotein gene (S gene). S gene was cloned into pMSCV_Neo vector (TakaRa Bio, USA) that vector have no GFP or luciferase reporter gene. After preparation of S gene contain plasmid p20012 (supplementary figure 2D), co-transfection was performed into viral production cell. pMD2G and pSPAX2 (Genemedi, China) packaging plasmid were used for retro based pseudovirus preparation. 9 × 10^6^ cells were seeded in a 75 cm^2^ tissue culture treated flask and cultured overnight. Then co-transfection was performed using Lipofectamine 3000 reagent as according to manufacturer’s protocol. After 6 hours of incubation, media was replaced with complete DMEM media. After 48 hours, media was collected and store it 4 °C. Additional 12 mL media was added into the flask and next day media was collected and combined with previously stored media. Then concentration and purification were performed by ultracentrifugation.^24^ After titer determination, pseudoviruses were stored at −86 °C freezer (ThermoFisher, USA).

#### In-vitro neutralization

ACE2 overexpressing HEK293 cell (Innoprot, Spain) were seeded in a two 96 well TC treated plate at a concentration of 2.2 × 10^4^ cells per well and overnight incubation was performed. One plate for adeno based pseudovirus and other plate for retro based pseudovirus. Two separate plate were used for serum preparation. Different rows of the plate were used for different group, such as A1-A10 for treatment group, B1-B10 for placebo, C1-C10, D1-D10 E1-E10 and F1-F10 for control, CR3022, commercial anti spike and only cell group. High concentration of CR3022, in-house developed, was used in these experiments. Sera from different mice of same group were collected and pool these sera for neutralization assay. 10 µL sera from vaccinated mice was added in 90 µL complete DMEM media. Then the serum was 2-fold serially diluted in complete DMEM media. For serum collected from different mice group, initial dilution was 10-fold with nine times 2-fold dilution. After completion of the serum dilution, 1.2 × 10^5^ pseudovirus in 50 µL was added into different wells that contained serially diluted serum and mixed properly. The SARS-CoV-2 pseudovirus and serum mixture was incubated for 1.5 hour at 37 °C. After incubation, 100 µL of pseudovirus and serum mixture was transferred on pre seeded cells. 5 µg/mL poly L-lysine (Wako, Japan) was added into each well for enhancing the transduction. Then, incubation was performed at 37 °C for 48 hours and after that readings for GFP fluorescence intensity were taken using Varioskan LUX (ThermoFisher, USA) machine. Number of virus particle inside the cells were determined by qPCR. After fluorescence analysis, media was removed and collected cells. Then heat inactivation was performed at 56 °C for 30 minutes. Cell was lysed and qPCR performed according to SYBR Green technology. In these experiment five wells from each group were selected and analyzed.

For retro based neutralization assay, qPCR was used to analyzed the copy number of S gene that integrated into cell. Copy number of S gene indicated the entry of pseudovirus into cell. Genomic DNA was extracted by MagMAX Express-96 Standard (ThermoFisher, USA) using Magmax DNA multi-sample ultra-kit. (ThermoFisher, USA). These genomic DNA was used for determination of S gene copy number by qPCR.

#### In-vivo neutralization

A total number of 18 albino male mice of 6-8 weeks were selected and isolated for the analysis. These mice were divided into 6 groups, 1 control and 5 treatment, comprising of 3 male mice in each group. The control group mice were immunized with 50 µL of placebo and treatment group mice were immunized with 1 µg/50 µL of BANCOVID vaccine. GFP Pseudotyped SARS-CoV-2 adeno virus (or treated as indicated in Figure: 6) were sprayed in the nasopharynx on 21-day post immunization. Nasopharynx and lung aspirate samples from mice were collected and analyzed for virus copy number using qPCR at indicated time point. Animals were sacrificed and lung section was performed and microscopic slides were prepared for fluorescence imaging (GFP) to detect virus load.

### Analysis

#### mRNA amplification

Purified mRNA, capped mRNA (vaccine candidate API), formulated LNPs and RNase treated formulated LNPs samples were used for mRNA amplification. RT-qPCR technique was performed according to GoTaq^®^1-Step RT-qPCR (Promega, USA) kit instructions. Primers, 0751F and 0752R, used are shown in supplementary table 3. Reverse transcription was done at 37 °C for 15 minutes then hold for 10 minutes at 95 °C for reverse transcriptase inactivation and GoTaq^®^ DNA Polymerase activation. Denaturation was done at 95 °C for 10 seconds, annealing at 44 °C for 30 seconds, extension at 68 °C for 30 seconds for 40 cycles. After completion of PCR cycle, melt curve was done for sample integrity checking.

#### mRNA identification

Capped mRNA, purified mRNA, formulated LNPs and formulated LNPs, treated with RNase samples, were analyzed by size exclusion chromatography (SEC). SEC was performed in Ultimate 3000 (ThermoFisher, USA) system using 10 mM Disodium hydrogen phosphate (Wako, Japan), 10 mM Sodium dihydrogen phosphate (Wako, Japan), 100 mM Sodium chloride (Merck, Germany), pH 6.6 as mobile phase. Biobasic SEC-300 (300 x 7.8 mm, particle size; 5 µm, ThermoFisher, USA) column was used with 1.0 mL/minute flow rate, 260 nm wavelength, 10 µL sample injection volume for 20 minutes.

### Humoral immunogenicity

#### Titer Analysis by ELISA

Serum from the mice of different groups were analyzed by enzyme-linked immunosorbent assay (ELISA) to determine sera antibody titers. ELISA plate (Corning, USA) was coated with 1µg/mL SARS-CoV-2 Spike S1+S2 ECD-His recombinant protein (Sino Biological, China) in Dulbecco’s phosphate-buffered saline (DPBS) (ThermoFisher, USA) for 2 hours at room temperature. Plate was washed for three time with DPBS + 0.05 % Tween 20 (Scharlau, Spain) and then blocked with PBS + 1 % BSA (ThermoFisher, USA) + 0.050 % Tween-20 for 2 hours at 37 °C. The plate was washed for 3 times then incubated with mouse sera and SARS-CoV-2 Spike antibody (Sino Biological, China) for 2 hours at 37 °C. After washing for 3 times, the plate was again incubated with Goat anti-Mouse IgG (H+L) Secondary Antibody, HRP conjugate (ThermoFisher, USA) for 50 minutes at room temperature. Final washing was done for 3 times and then developed the colorimetric reaction with Pierce TMB substrate (ThermoFisher, USA) for 10 minutes. The reaction was stopped with 1N hydrochloric acid (HCl) and the plate was read at 450 nm wavelength within 30 minutes.

#### Isotyping analysis by ELISA

For isotype analysis, Pierce Rapid ELISA Mouse mAb Isotyping kit (ThermoFisher, USA) was used. Serum samples from 4 subjects of treatment 2 and 3 were analyzed. All the steps were followed as per supplier’s instructions.

#### Antibody binding affinity by SPR

The BIAcore T200 equipment (GE Healthcare, USA) and Amine coupling kit (GE Healthcare, USA) were used for immobilization of SARS-CoV-2 Spike S1+S2 ECD-His recombinant protein (Sino Biological, China) in Series S Sensor Chips CM5 (GE Healthcare, USA). First the flow cell surface of Series S Sensor Chips CM5, was activated by injecting a mixture of EDC/NHS (1:1) for 7 minutes. Then 70 µL of 50 µg/mL S1+S2 protein was prepared in sodium acetate at pH 5.0 and injected over the activated surface at 10 µL/min flow rate. Residual NHS-esters were deactivated by a 70 µL injection of 1 M ethanolamine, pH 8.5. The immobilization procedure was performed by using running buffer HBS-EP, pH 7.4 (GE Healthcare, USA).

5 samples containing 1 µL mouse serum each were analyzed using surface plasmon resonance (SPR) technology to analyze the binding affinity of the antibody pool. The first, second, third, fourth and fifth samples were 1 µL of pre-immune mouse serum, 1 µL of MabSelect resin (GE Healthcare, USA) pulldown mice serum after 14 days of immunization, 1 µL resin pulldown mice serum with 0.5 µL of 500 µg/mL of SARS-CoV-2 Spike S1+S2 ECD-His recombinant protein, 1 µL of resin pulldown mice serum with 0.5 µL of 500 µg/mL SARS-CoV-2 Spike RBD-His recombinant protein, and 1 µL of resin pulldown mice serum with 0.5 µL of 500 µg/mL SARS-CoV-2 Spike S2 ECD-His recombinant protein, respectively. These samples were flown through over the active flow cell surface of CM5 chip for binding analysis. Glycine-HCl of pH 2.5 was used for regeneration. All samples were diluted in 1 x HBS-EP at pH, 7.4 running buffer.

### Cellular immunogenicity

#### SARS-CoV-2 surface glycoprotein peptide mapping

SARS-CoV-2 Spike S1+S2 ECD His recombinant protein (Sino Biological, China), S2 ECD-His Recombinant Protein (Sino Biological, China), and RBD-His Recombinant Protein (Sino Biological, China) were digested and purified according to ThermoFisher Pierce Trypsin Protease, MS grade instructions (supplementary method 2). 1 µg of digested peptides were loaded into mass spectrometry system (Q Exactive Hybrid Quadrupole-Orbitrap MS manufactured by ThermoFisher Scientific, USA). For separation of peptides Hypersil gold C18 (100x2.1 mm; particle size: 1.9 µm, ThermoFisher, USA) column was used. Column oven temperature was set at 40 °C and eluted in 5 – 40 % mobile phase B (0.1 % formic acid in acetonitrile) and 95– 60 % mobile phase A (0.1% formic acid in water) gradient with 0.300 mL/min flow rate for 65 minutes. Peptide elution were checked by UV absorbance at 214 nm. For peptide identification, data dependent mass spectrometry was performed where full-MS scan range was 350 m/z to 2200 m/z, resolution was 70,000, AGC target was 3E6, maximum IT was 100 milliseconds (ms), and data dependent mass spectrometry resolution was 17,500, AGC target was 1E5, maximum IT was100 ms. After getting raw data from mass spectrometry system, data analysis was performed in BioPharma Finder (ThermoFisher, USA) using variable parameters to get confident data, and then data were combined in one map to visualize complete fragmentation (supplementary figure 6).

**Figure 6:**
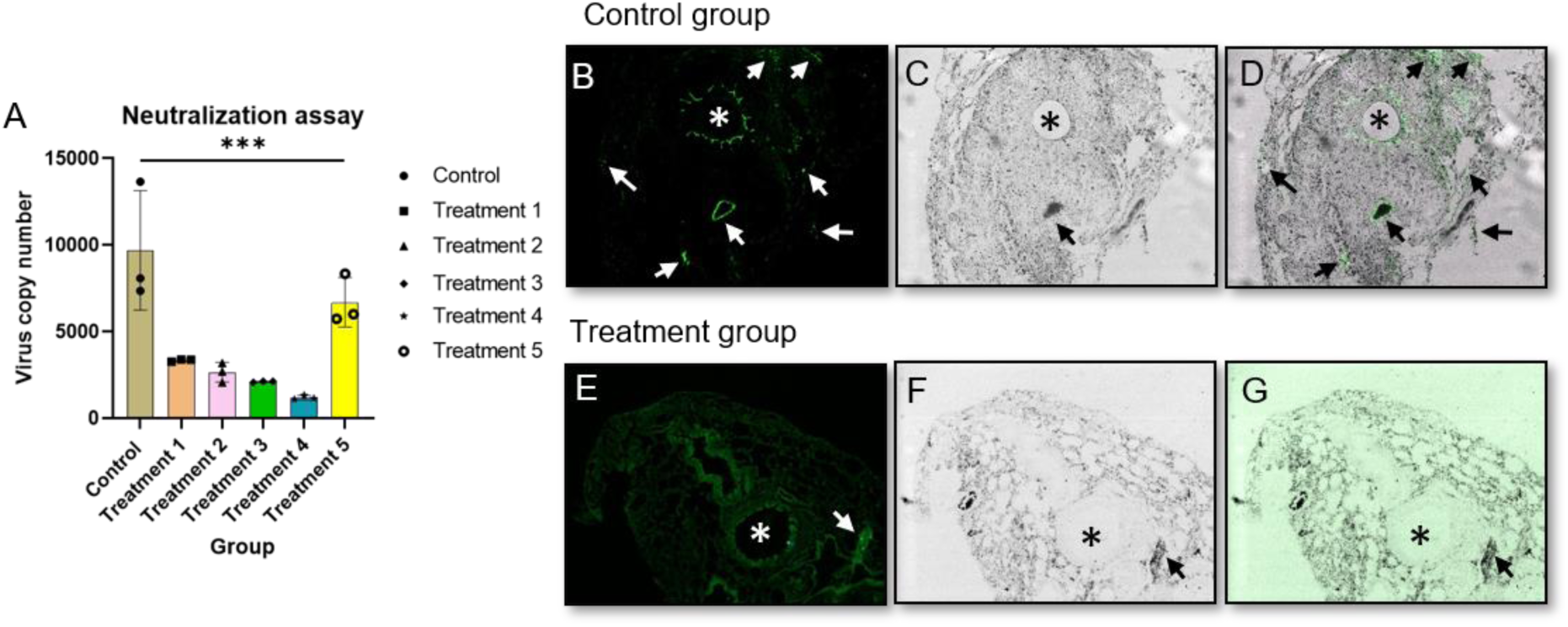
*In-vivo* neutralization assay, lung section, * indicates trachea and arrow indicate infection. (A) (B) fluorescence image of lung section of control group mouse, (C) trans image of lung section of control group mouse, (D) overlay image of lung section of control group mouse, (E) fluorescence image of lung section of treatment group mouse, (F) trans image of lung section of treatment group mouse, (G) overlay image of lung section of treatment group mouse, intentional green color enhancement was done to observe any GFP intensity for panel G.

On the other hand, full length SARS-CoV-2 surface glycoprotein was digested computationally (ExPASy PeptideMass: https://web.expasy.org/peptide_mass/) via trypsin (supplementary figure 7).

#### Mouse splenocyte isolation, peptide stimulation and Flow cytometric analysis of T cell (CD4^+^) populations

Male and female BALB/c swiss albino mice were sacrificed and splenocyte were harvested using in-house developed method (supplementary method 3). Harvested splenocyte were RBC lysed and cultured at 37°C and 5% CO2using RPMI complete media where sacrificed mice sera were used instead of fatal bovine serum (FBS). A time-lapse video at 40X magnification for splenocyte was captured after 24 hours of culture. Isolated splenocytes from mice were either stimulated with S-peptide pool or buffer. After 6 hours, media were collected, cells were washed twice with PBS and incubated for 12 hours. This samples were considered as 18 hours’ sample. Samples were collected again for cytokine secretion assay and cells were processed for Flow cytometric analysis of T cell populations. Intracellular cytokine staining of cells were stained with following antibodies with maintaining supplier’s instructions: V500 anti-mouse CD45 (BD Bioscience, USA), FITC anti-mouse CD4 (ThermoFisher, USA), anti-mouse IL-2 (ThermoFisher, USA), Alexa Fluor® 594 conjugate secondary antibody (ThermoFisher, USA), anti-mouse IL-6 (ThermoFisher, USA), Alexa Fluor® 594 conjugate secondary antibody (ThermoFisher, USA), in-house developed TNF alpha fusion protein, anti Fc primary antibody (ThermoFisher, USA), Alexa Fluor® 594 conjugate secondary antibody (ThermoFisher, USA) and no live/dead staining was used. Cells were washed, fixed, stained and stored at 4 °C. After 48 hours, cell events were acquired using an FACS Lyric (BD Biosciences), followed by FlowJo software (FlowJo LLC, Ashland, OR) analysis (supplementary figure 8, 9, 10).

#### IL-2 and Il-6 titer

ELISA plate (Corning) was coated with 1µg/mL IL-2 polyclonal antibody (ThermoFisher, USA) in Dulbecco’s phosphate-buffered saline (DPBS) (ThermoFisher, USA) for 2 hours at room temperature. After coating, Plate was washed for 3 times with DPBS + 0.05 % Tween 20 (Scharlau, Spain) and then blocked with PBS + 1 % BSA (ThermoFisher, USA) + 0.050 % Tween 20 for 2 hours at 37 °C. After blocking, Plate was washed for 3 times and incubated with IL-2 and mouse splenocyte samples for 2 hours at 37 °C. Plate was then washed again and incubated with IL-2 monoclonal antibody (ThermoFisher, USA) for 2 hours at 37 °C. After washing for 3 times, the plate was again incubated with Goat anti-Mouse IgG (H+L) Secondary Antibody, HRP conjugate (ThermoFisher, USA) for 50 min at room temperature. Final washing was done for 3 times and then developed with Pierce TMB substrate (ThermoFisher, USA) for 10 min and then stop with 1N hydrochloric acid (HCl). Finally, plate was read at 450 nm wavelength within 30 min of stopping reaction. Titers were quantified through 5 parameter logistics best fit curve.

For IL-6 analysis, IL-6 Mouse ELISA kit (ThermoFisher, USA) was used. All the steps were performed as per manufacturer instructions.

## Results

### Bioinformatics analysis to initiate the designing of ‘BANCOVID’

We have started with alignment of available sequences of SARS-CoV-2 spike (S) protein. In march, 2020, we found total 15 D614G sequences out of 170 reference sequences of SARS-CoV-2 (Supplementary Figure: 1A). The full sequence alignment is given elsewhere (data not shown). Hydropathy profile showed a minor variation in relevant protein between D614 and G614 genotypes (Supplementary Figure: 1B and C). Relevant 3D modeling suggested that there might be higher angular strain on G614 than the D614, which could affect the stability and atomic distance with the neighboring atoms (Figure: 1E and F). Our observation has been recently validated by others. ^14, 25, 26^

### Construction, antigen expression and formulation of ‘BANCOVID’

We have obtained the ORF for the SARS-CoV-2 spike with G614-translating codon from a clinically confirmed COVID-19 patient through PCR amplification (Accession No.: MT676411.1). Necessary modifications were performed to obtain the desired clone in pET31b vector as described in ‘Materials and Method’ section. The schematic diagram of the target gene and construction scheme are shown in Figure: 1A and Supplementary Figure: 2A and B, respectively. The *in vitro* transcription (IVT) process was modified to obtain high yield and desired quality of mRNA (Figure: 1B). We have obtained the capped-mRNA with a 130-nucleotide residue-long poly A tail. The mRNA sequence with poly A tail was confirmed by DNA sequencing after reverse transcription (Figure: 1C); Accession No.: MWO45214. The IVT process was tuned to obtain desired mRNA with high yield and quality (Figure: 1C). The mRNA was encapsuled in lipid nano particle (LNP) ranging from 60 – 140 nm with the final pH of 7.2. We did a pilot study with limited numbers of mice to identify the suitable mRNA-LNP size for our formulation. mRNA-LNP either smaller than 70 nm or larger than 110 nm did not generate considerable immunological response even with a dose of 10 ng/mice (data not shown). To obtain the best process control for the dose production, we therefore, set our mRNA-LNP size range at 85±10 nm. We used mRNA-LNP of this range throughout the rest of the experiments (Figure: 1E). LNP without SARS-CoV-2 S-mRNA was used as placebo.

### Local tolerance and toxicity

Control, treatment and placebo group comprising 3 male mice each were used for local tolerance testing. Pictures of the site of injection before and 24 hours after injection are shown in Figure 2A (top and bottom panels, respectively). No detrimental physical consequences of administration were observed such as, local trauma following injection and/or physicochemical actions of the vaccine from local toxicological or pharmacodynamics effects. No sign of erythema or erythredema were observed in muscle tissue section from the site of injection (Figure: 2B). Complete blood count (CBC) count from different groups indicated good health status of mice; all parameters were in normal physiological range (Figure: 2C – J). There were no signs for anemia, infection, inflammation, and bleeding disorder. Liver function tests (LFTs) such as alanine transaminase (ALT) and aspartate aminotransferase (AST) were performed to confirm clinical suspicion of potential liver injury or disease and to distinguish between hepatocellular damage and cholestasis (Figure: 2K and L). Blood urea nitrogen (BUN) was tested to evaluate the health of kidneys, such as kidney disease or damage (Figure: 2M). Data for ALT, AST and BUN were in normal range and no significant changes were observed between pre-immunization and after immunization.

**Figure 1:**
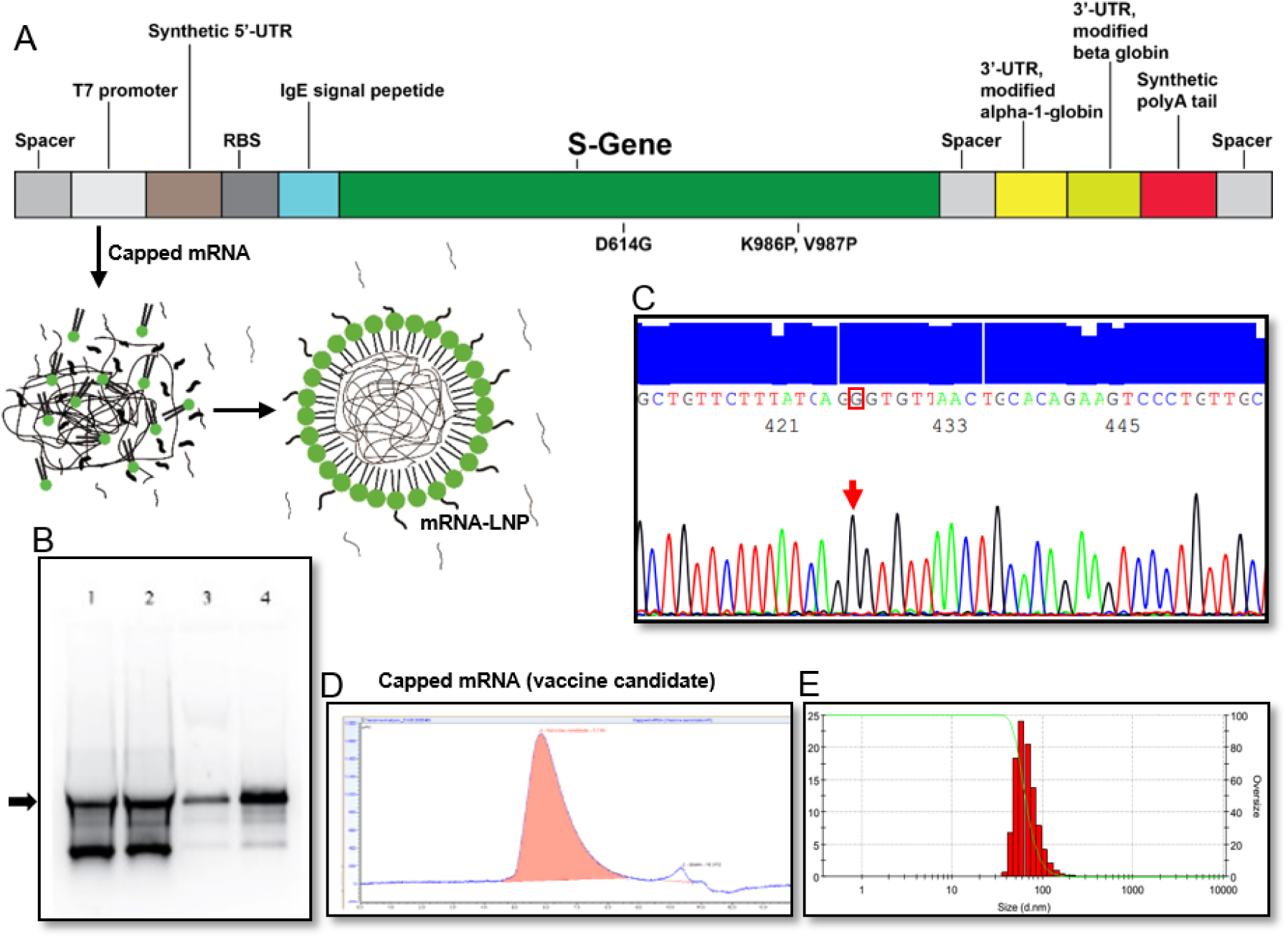
Target construction, amplification, IVT optimization, purification, and LNP formation. (A) Graphical representation of linear DNA construct for mRNA transcription, (B) DNA sequencing electropherogram data of D614G sequence in the target, (C) IVT optimization where Lane 4 is the optimized condition, (D) Identification of purified capped mRNA by SEC-HPLC, (E) size distribution of mRNA-LNP dose formulation.

**Figure 2:**
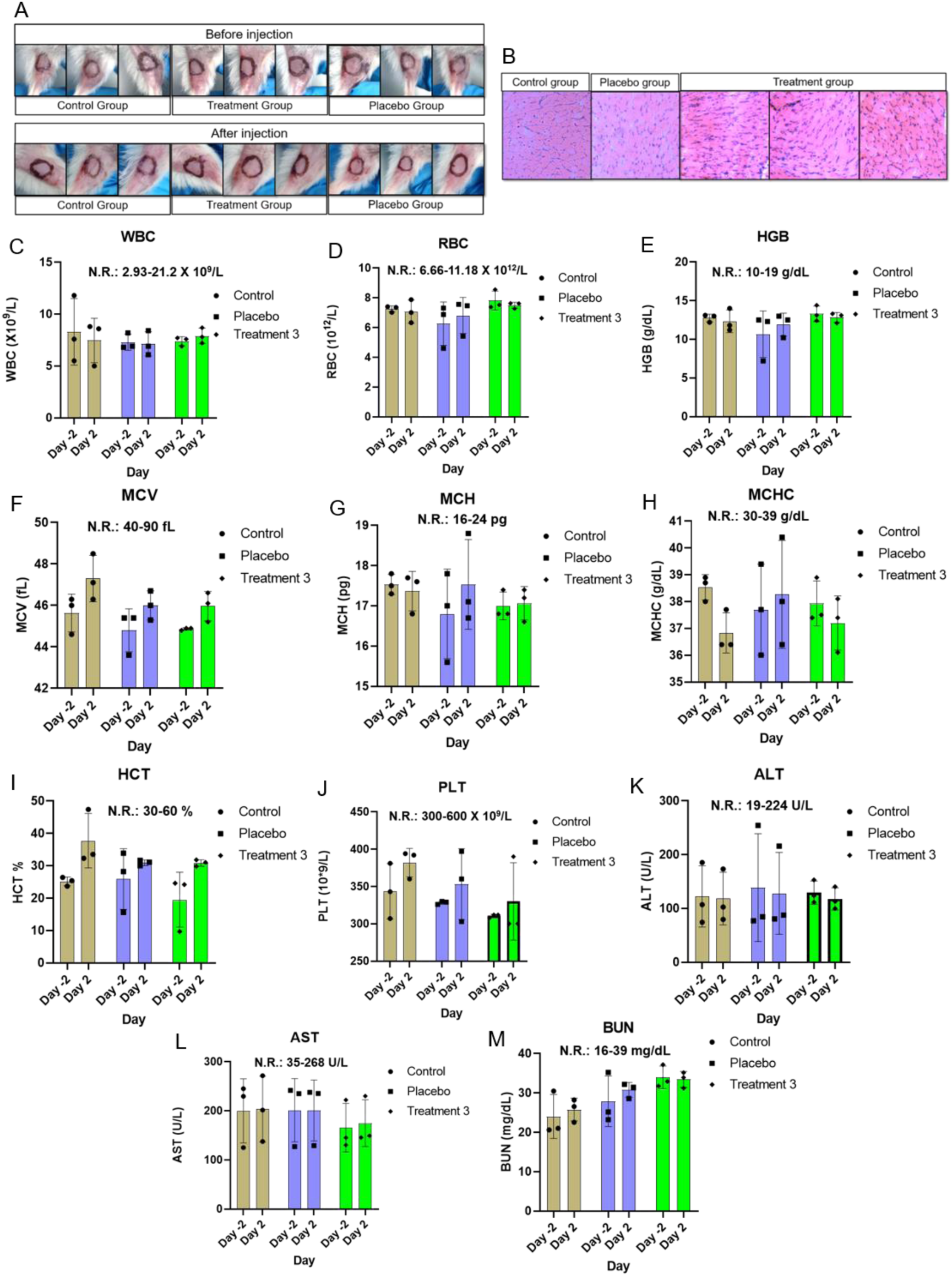
Local tolerance and CBC analysis. (A) check for sign of visible adverse reaction of administration before and after injection, (B) HE stained tissue from site of injection for erythema and edema, (C) WBC, white blood count, (D) RBC, red blood cell, (E) HGB, hemoglobin, (F) MCV, mean corpuscular volume, (G) MCH, mean corpuscular hemoglobin, (H) MCHC, mean corpuscular hemoglobin concentration, (I) HCT, hematocrit, (J) PLT, platelet, (K) ALT/GPT, alanine transaminase, (L) AST/GOT, aspartate aminotransferase, (M) BUN, blood urea nitrogen.

### ‘BANCOVID’ induces high and Th-1 biased antibodies against full-length SARS-CoV-2 S-protein

Immunization of mice with mRNA-LNP produced specific titer at a dose dependent manner (Figure: 3A). Low dose (0.1 µg/mice) immunization produced moderate level of antibody response (Figure: 3A, Treatment 1). We found best antibody response with 1 µg/mice dose (Figure: 3A, Treatment 2). High dose (10 µg/mice) immunization produced higher level of titer but the response among the mice were inconsistent (Figure: 3A, Treatment 3). The subtyping analysis revealed that the titer contains balanced ratio of IgG2a and IgG1 in 7-day post immunization sera, and it remains stable for 14-day post immunization sera (Figure: 3B, Treatment 2). Similar trend was observed for (IgG2a + IgG2b) and (IgG1+IgG3) (Figure: 3C, Treatment 2), which has suggested that the antigenic response was CD4+Th1-biased. High dose (10 µg/mice) injected mice sera also produced similar response (Figure: 3B and C, Treatment 3). The complete isotyping data is shown in Supplementary Figure: 5. To check whether the immunization have generated antibody pool spanning for the whole antigen or for any specific domain (S1 or S2), we have chosen surface plasmon resonance (SPR) experiment. The S protein chip recognized high-affinity antibody from the anti-sera (Figure: 3D). The response was attenuated significantly for S-protein(s) (S, S1 and S2) pretreated sera (Figure 3D). S and S1 pretreatment showed similar and strong inhibitory response while S2 pretreatment showed comparatively moderate inhibitory response. The purified Ig from the pooled anti-sera produced significantly pronounced response (Figure: 3E). The SPR data clearly showed that the vaccination has produced specific antibody pool against the full-length of S protein.

**Figure 3:**
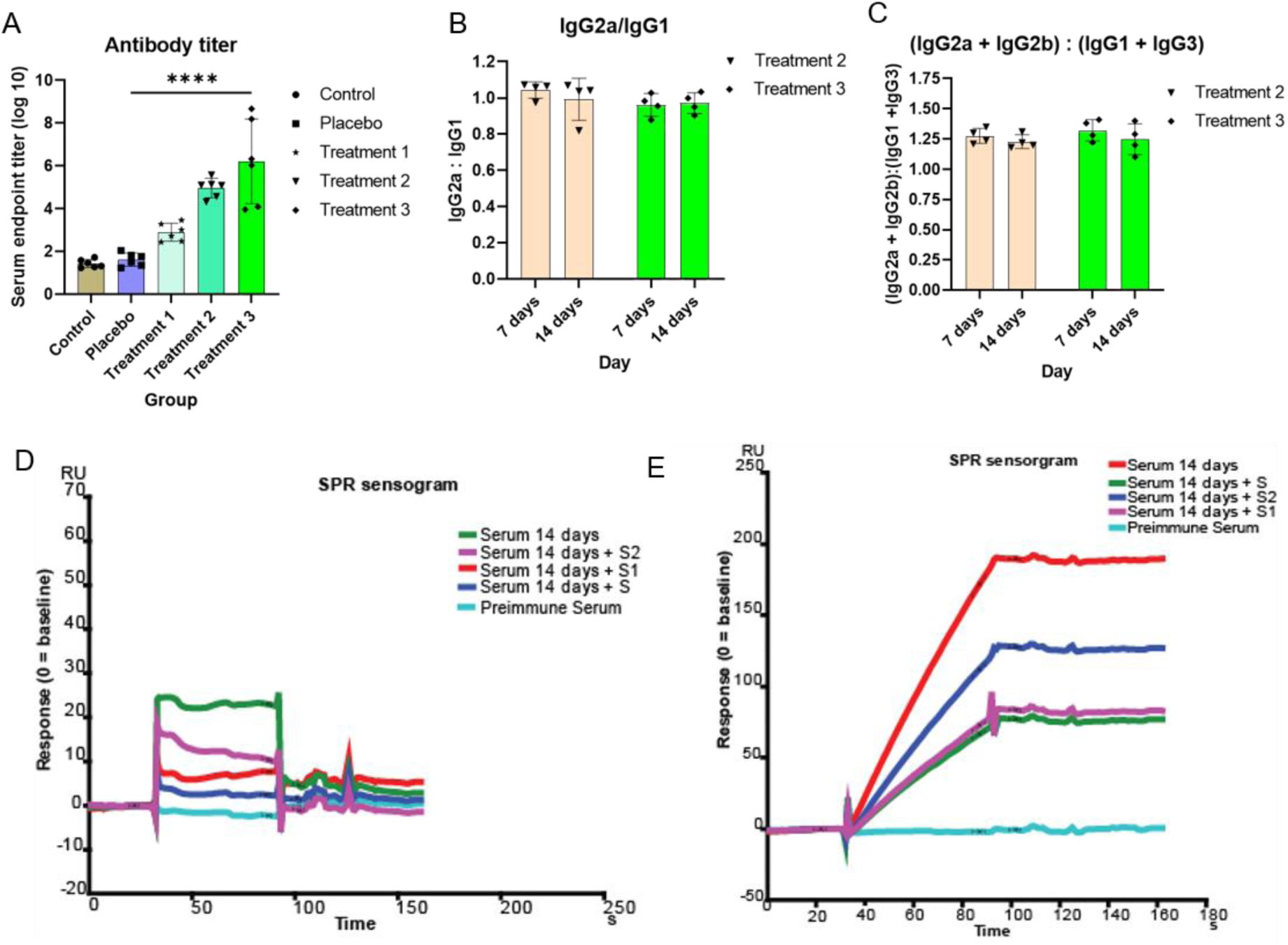
Antibody affinity and titer analysis. (A) antibody titer analysis from serum of different groups after 14 days of immunization, all the group data were compared (1.0 µg and 10.0 µg) by Mann-Whitney test, ****= p-value< 0.0001, (B) ratio of IgG2a and IgG1 in treatment 2 and treatment 3 group, (C) ratio of IgG2a+IgG2b and IgG1+IgG3 in treatment 2 and treatment 3 group. (D) serum antibody affinity analysis, (E) resin pull-down serum antibody affinity analysis.

**Figure 4:**
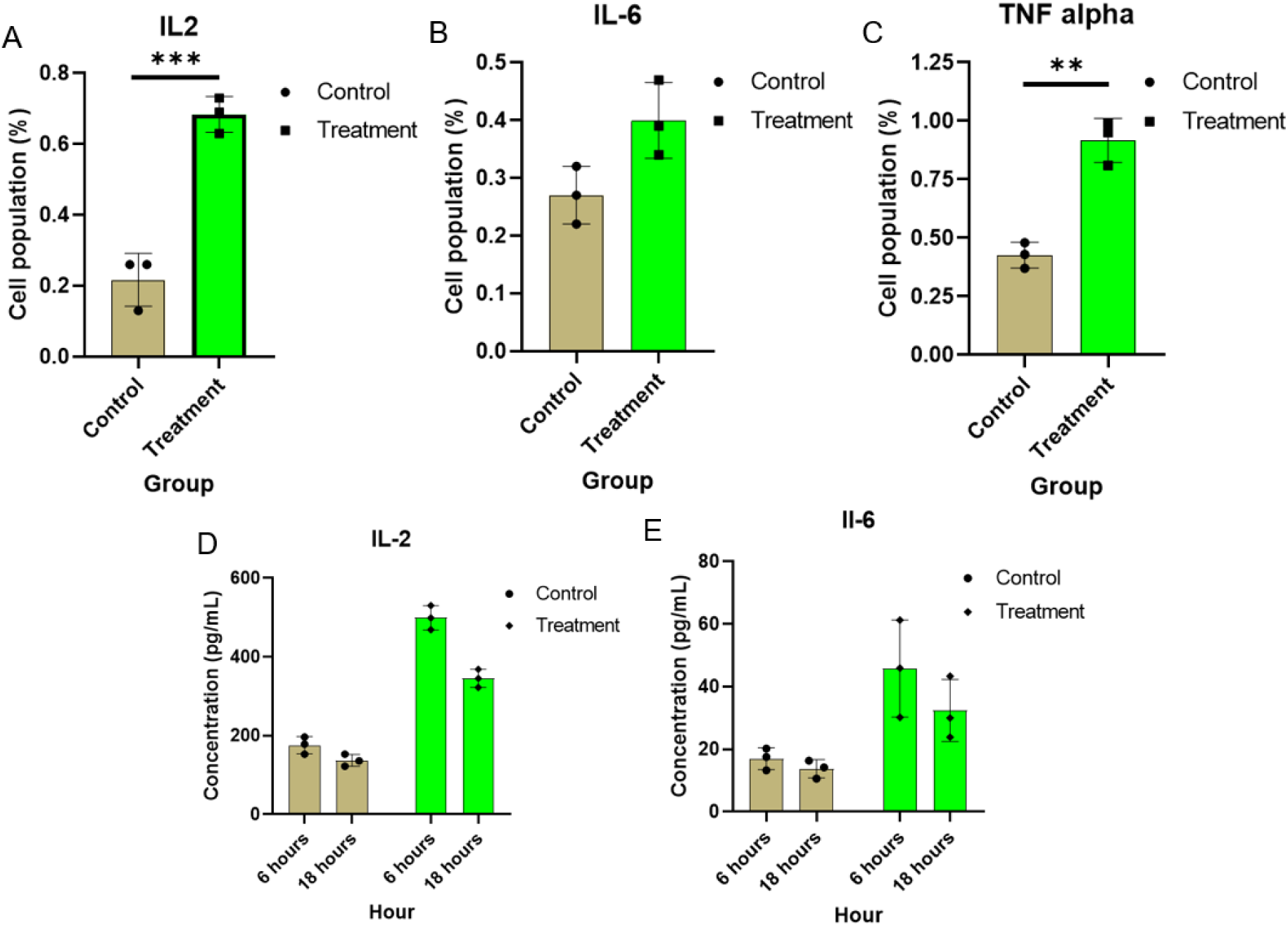
Cellular immune response analysis (internal and secretory cytokine) in control and treatment group; unpaired T-test were performed between control and treatment groups; ***= p-value<0.001, **= p-value<0.01, (A) IL-2 expressing cell population percentage of control and treatment group, (B) IL-6 expressing cell population percentage of control and treatment group, statistically non-significant, (C) TNF-α expressing cell population percentage of control and treatment group, (D) secretory IL-2 concentration analysis between control and treatment groups at 6 and 18 hours, (E) secretory IL-6 concentration analysis between control and treatment groups at 6 and 18 hours.

**Figure 5:**
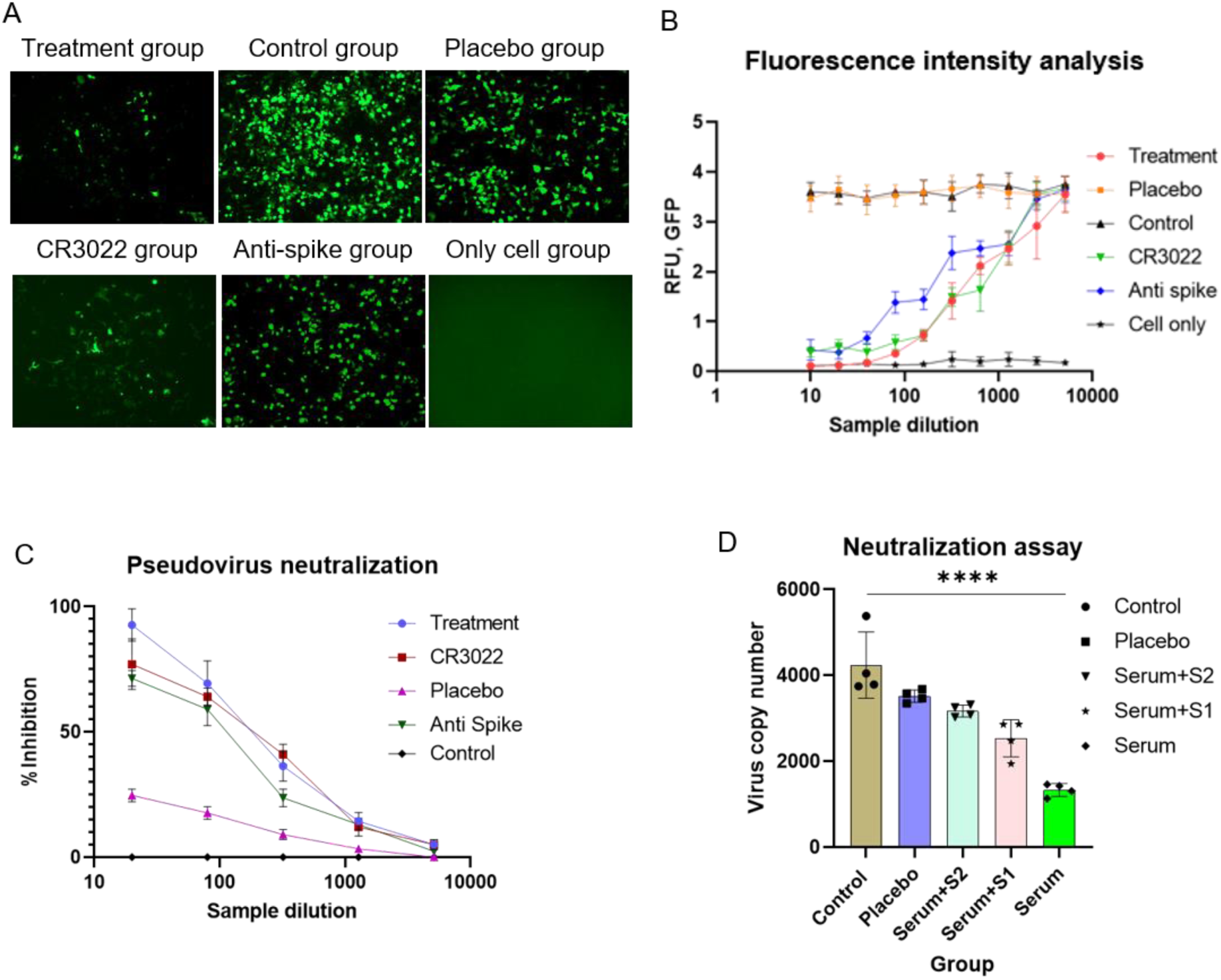
*In-vitro* neutralization assay. (A) Image of Green fluorescence protein (GFP) expression after adeno-based SARS-CoV-2 pseudovirus neutralization assay from 2^-4^ sample dilution, (B) correlation between SARS-CoV-2 antibody from mice sera and intensity of GFP in different experimental group. For treatment group with the decrease of the antibody concentration, the intensity of GFP expression increased, which indicated the inhibition of SARS-Cov-2 pseudovirus into ACE2 overexpressed HEK293 cell (ACE2-HEK293 cell), (C) adeno-based SARS-CoV-2 pseudovirus neutralization percentage at different sample dilution, (D) HIV-1 based SARS-CoV-2 pseudovirus copy number analysis by qPCR; all the samples were compared by one-way ANNOVA method, ****= p-value< 0.0001.

### Cellular and cytokine responses to ‘BANCOVID’

We further characterized the cellular response and induction of specific cytokines in response to vaccination. The splenocytes obtained from vaccinated mice were re-stimulated with a library of SARS-CoV-2-S peptide pool. The stimulated splenocytes generated significantly higher population of CD4+Th1 cytokine IL2- and TNFα-expressing cells (0.68 ± 0.05 and 0.9 ± 0.09, respectively) compare to the placebo treated group (0.22 ± 0.08 and 0.43 ± 0.06, respectively) (Figure: 4A and B). CD4+Th2 cytokine IL6-expressing cells were moderately increased in stimulated splenocytes of vaccinated mice compare to the placebo-injected mice (0.40 ± 0.07 and 0.27 ± 0.05, respectively) (Figure: 4C). The IL2 and IL6 secretion from restimulated splenocytes were found significantly increased for vaccinated group (499.10 ± 30.80 pg/ml and 45.78 ± 15.52 pg/ml, respectively) over the placebo group (175.71 ± 21.92 pg/ml and 16.96 ± 3.53 pg/ml, respectively) (Figure: 4D and E). Sustained secretion of cytokines from splenocytes of vaccinated mice were observed even after 12 hours of withdrawal of the S-pool protein stimulation; (345.17 ± 22.85 pg/ml of IL2 and 136.87 ± 15.18 pg/ml of IL6, respectively) (Figure: 4D and E). Higher level of sustained Th1 specific cytokine response over Th2 specific cytokine suggested a stable and balanced Th1-biased immunologic response after administration of ‘BANCOVID’ vaccine.

### ‘BANCOVID’ induces neutralization of SARS-CoV-2-S pseudo-type viruses

#### In-vitro neutralization

Sera of vaccinated mice inhibited infection of GFP-expressing pseudo-type SARS-CoV-2-S adenovirus in hACE2-expressing HEK293 (ACE2-HEK293) cell in a dose dependent manner (Figure: 5A and B). Neutralization assay demonstrated that there is a correlation between the intensity of GFP and SARS-CoV-2 specific antibody for vaccinated mice. Higher concentration of SARS-CoV-2 antibody efficiently neutralize the entry of the pseudovirus into the ACE2-HEK293 cell. IC50 value for GFP-inhibition were found significantly higher for the anti-sera (∼3 µg/mL) compared to the CR3022 and a commercially available polyclonal mouse antibody against S-protein (∼7 µg/ml). To confirm whether the GFP signal was generated from GFP shedding from degraded virus instead of functional virion, we have analyzed the viral copy number using real-time PCR. The data showed correlation with the GFP and gene copy analysis (Figure: 5D). HIV1-based SARS-CoV-2-S pseudo type virus infection was also significantly inhibited by 1 µg/mice dose anti-sera compared to the placebo anti-sera (Figure: 5D, Serum). Either S1 or S2 protein pretreatment nullified the inhibition capacity of anti-sera (Figure: 5D, Serum+S1 and Serum+S2) confirmed that the inhibition property for the HIV1-based SARS-CoV-2-S pseudo type virus is related to the vaccination.

#### In-vivo neutralization

Next, we attempted to check whether immunization can protect mice from GFP-expressing pseudo-type adenovirus. Virus were sprayed through nasopharyngeal space of mice either in buffer or pretreated with immunized sera. The anti-sera-treated SARS-CoV-2-S adenovirus produced lower copy of virus compared to the buffer-treated virus (Figure: 6A, Treatment 2 and Treatment 1, respectively). The copy number of virus was found reduced further from day 2 to day 3 (Figure: 6A, Treatment 4 and Treatment 3, respectively). These data clearly revealed that though the anti-serum has significant inhibitory capacity against viral infection but systemic immune-protection is essential for better protection. Lower copy number of virus over the time indicated that the cellular immunity is also necessary, along with humoral immunity, for viral clearance. The anti-sera treated with S1+S2 protein failed to inhibit SARS-CoV-2-S adenovirus infection in the placebo-injected mice (Figure: 6A, Treatment 5), and proved that the inhibition and neutralization of the SARS-CoV-2-S pseudo-type virus is correlated with the immunogenic response generated due to ‘BANCOVID’.

## Discussion

The G614 variant was first identified in China and Germany in January, 2020. It was a rare genotype before March, 2020, which then quickly became the major circulating genotype worldwide.^14^ Cardozo *et. al.*, reported in May, 2020 that G614 genotype is associated with increased case fatality rate over D614.^27^ Scientific findings evidently demonstrated that the G614 variant is ∼10 times more infectious over the D614 genotype.^28, 29^ It has been revealed from *in vitro* and clinical data that G614 variant has a distinct phenotype, and there likely be a huge impact of this mutation on infection, transmission, disease onset, disease prognosis, as well as on vaccine and therapeutic development.^14, 30, 31^ We found the sequence G614 from a PCR-confirmed patient in May (Accession No. MT676411.1). Based on the scientific information available then we have predicted that this variant may become dominant in future, and at that period of time there was no information for any vaccine candidate in development considering G614 genotype. We, therefore, decided to develop the vaccine considering this mutation.

Since the vaccine may banish (BAN) COVID-19 (COV) and originated from Bangladesh (BAN) we therefore named it ‘BANCOVID’. The design consideration for the vaccine was to obtain high-expressing spike protein as antigen in a putative perfusion stabilized condition. Comparative design features of ‘BANCOVID’ with the available published information for relevant vaccine candidates in development are shown in Table: 1. ‘BANCOVID’ mRNA has few features along with the G614-targetted mutations, which are different than the other relevant known candidates. Ribosome binding site, IgE-signal sequence by replacing the native 13 amino acids from the N-terminal of the SARS-CoV-2 S protein, 3’ UTR constituted with the 3’UTRs of alpha and beta globin gene in tandem are worth to mention. We have used 70 – 100 nm LNP to deliver the mRNA. LNPs out of this range did not elicit considerable antibody response in a separate pilot study (data not shown); the best range was observed with 85±10 nm of LNP. The LNP-sizes determine the delivery efficiency of the cargo to the target cells.^32^ The pH (7.2) of our formulation buffer for mRNA-LNP is also lower than the other relevant references (7.4∼8.0).^2,7,8,9^ Lower pH helps quick release of the cargo from endosomal compartment and protects mRNA from acid hydrolysis and lysosomal digestion in intracellular milieu.^33^ Together, numbers of minute changes in the design context likely playing in concert and produced quick, balanced, stable Th1-IgG2-biased antibody response.

‘BANCOVID’ immunization did not produce any noticeable effect for local or systemic toxicity as primarily evident by the absence of four cardinal signs of inflammation: redness (Latin rubor), heat (calor), swelling (tumor), and pain (dolor). There was no erythema or erythredema as well in any injection site. The CBC and blood chemistry data did not show significant changes in relevant profiles and has been suggesting that the vaccine behaves safely in animal.

A balanced response between Th1 and Th2 is desired to achieve safe and effective humoral immunity performance.^35^ ‘BANCOVID’ has produced well-balanced IgG1 and IgG2 response by 7^th^ day postimmunization and remained similar on 14^th^ day postimmunization sera, which is suggesting a stable antibody response during the sampling period. Along with opsonizing characteristics, IgG2 has higher affinity to its receptors and have superior complement system activation potential over IgG1.^35, 36^ Accordingly, ‘BANCOVID’-mediated higher ratio of IgG2 than IgG1 has suggested that higher capacity of the antibody pool to clear antigen from the system. The ratio of IgG2a and IgG1, and cytokine-stained CD4^+^ and CD8^+^ T cell population showed a Th1-bias response. Since mouse IgG2 is equivalent to human IgG1^35, 36^ therefore, it is plausible that ‘BANCOVID’ will elicit effective cellular and humoral response against SARS-CoV-2 in human.

The early vaccine development initiatives were taken before the G614 variant became predominant. Therefore, there are no specific vaccine tested so far in human with G614 variant-related molecule. Studies with the sera obtained from COVID-19 patients showed variable results regarding the neutralization propensity of D614 and G614 genotype. With 88 sera from a high-incidence community, Sadtler *et al.*, showed that antibody pool did not differentiate between D614 and G614 binding.^15, 37^ However, a few data point stayed out of the correlation trend in their study, which might be linked with the functional variations associated with the SARS-CoV-2 variants. Korber *et al.*, with 6 convalescent sera, showed that D614 and G614 both types of VSV-pseudovirus can be inhibited by the sera; though G614 and D614 showed little variation in their responses to the assay^14^. They further showed that the G614 genotype produced higher titers against pseudoviruses from *in vitro* experiments. The variations observed in both of the studies might not be just a coincident rather suggesting potential differences in *modus operandi* between the G614 and D614 variants. The proposition is supported by the Huang *et al.*; they have showed that 7% of the convalescent sera out of their 70 samples failed to neutralize G614 variant of pseudovirus.^38^ All of these studies did not identify whether the subjects were infected by either D614 or G614 variant, which could have revealed better insight for the correlation of the observation.

The roles of G614 mutation on constitutive infection have been attributed to its conformational change. It has been proposed that the -COOH group of D614 forms hydrogen bond with the - OH group of T859 across the S1/S2 interface, which cannot form in G614.^14^ On the contrary, structural modeling studies revealed that “the D614G substitution creates a sticky packing defect in subunit S1, promoting its association with subunit S2 as a means to stabilize the structure of S1 within the S1/S2 complex.^30^ In other words, the D614G mutation in fact promotes the S1/S2 association and stabilize the spike.^30^ The finding is in accordance with the observation that G614 has a greater stability originating from less S1 domain shedding and greater accumulation of the intact S protein into the pseudovirion.^29^ It has also been reported that G614 mutation promoted the interaction of two of the three S glycoprotein chains with the RBD whereas only one chain from D614 interacts with RBD.^26^ This interaction creates favorable conformation of the RBD to its partners resulting higher access for effective binding and infection.

Predecessor SARS-CoV virus entry and infection was shown promoted by protease-mediated processing of the spike protein.^39^ It has been postulated that SARS-CoV-2 also likely be adopting such properties by acquiring G614 genotype by incorporating protease processing site.^40^ Consequently, it has been shown that indeed G614 protein has been cleaved by serine protease elastase-2 more efficiently than the D614 variant.^38^ They further showed that G614 pseudovirus infection of 293T-ACE2 was potentiated in the presence of elastase-2, which can be blocked by elastase inhibitor. These findings corroborate the fact that G614-targetted vaccine is necessary.

Two prominent antigenic sites on the S-protein have been proposed those are spanning 614 position: V515-D614 and D614-A647.^30, 41^ The role of V515-D614 domain is not known but the D614-A647 is a dehydron wrapped intramolecularly by residues D614, V615, T645, R646 and A647, and forms a salt bridge with D614. The salt bridge contributes to stabilize D614-A647 in the uncomplexed S1 and inhibits the S1/S2 association. G614 diminishes the salt bridge formation and S1/S2 association resulting interaction with the RBD to facilitate higher infection.^30^ Therefore, blocking of G614 with a specific antibody would inhibit such acquired fitness of SARS-CoV-2. ‘BANCOVID’ immunization has produced a pool of antibody that covers the whole length of the spike protein suggesting that highly likely relevant antibody-mix against these domains have been developed. Since relevant domains are highly glycosylated therefore, we could not obtain homotypic peptides for definitive characterization of the purified antibody pool against these predicted antigens. Further study will be attempted to reveal the relevant scientific aspects. Nevertheless, the findings clearly demonstrated that ‘BANCOVID’ is safe for *in vivo* administration, and elicits balanced and stable cellular and humoral response that neutralize SARS-CoV-2 spike protein-mediated infection. Currently, we have been preparing for the phase-1 clinical trial. Appropriate clinical trial will reveal further insight regarding the significance of G614-targetted vaccine for the efficient management of COVID-19 pandemic.

The recent metadata analysis on more than 5000 clinical samples revealed that 100% of the second-wave of infection has been associated with G614 variant, which is emphasizing the need for G614-targetted vaccine for managing this uncontrolled infection.^13^ Therefore, the rapid transition of the 1^st^ G614-targetted vaccine ‘BANCOVID’ to clinical trial would be highly rewarding.

## Acknowledgements

The study was funded by Globe Biotech Ltd. We thank Md. Harunur Rashid, the chairman of Globe Pharmaceuticals Group of Companies, Ahmed Hossain, Md. Mamunur Rashid and Md. Shahiduddin Alamgir, the directors of Globe Pharmaceuticals Group of Companies for their continuous support and encouragement. We also thank Md. Raihanul Hoque, Dibakor Paul, Biplob Biswas, Zahir Uddin Babor, Mithun Kumar Nag and Imran Hossain for their support for facility and information management system.

---

**Supplementary figure 1:**
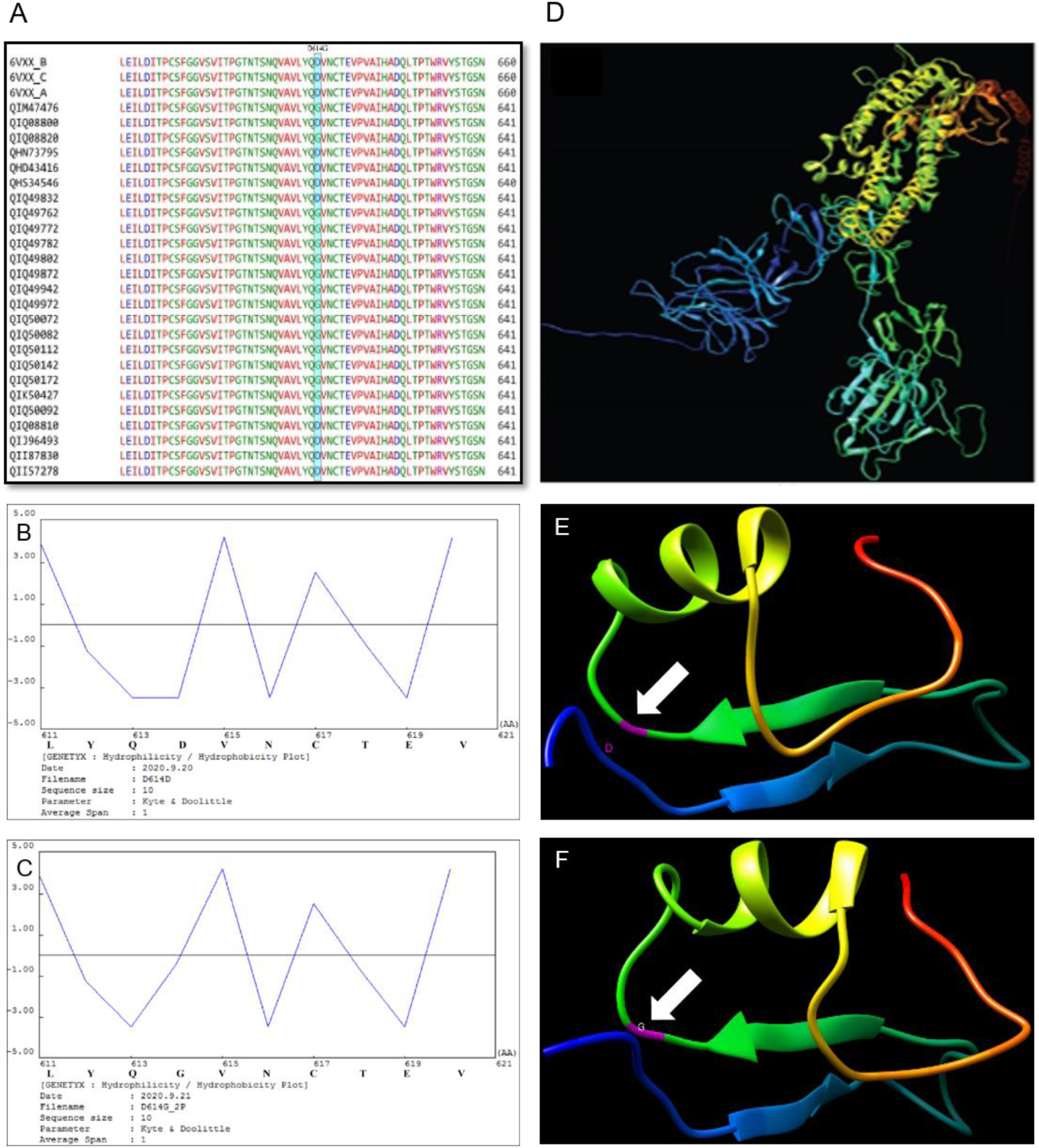
Target gene selection and modification; (A) Sequence alignment of SARS-CoV-2 surface glycoprotein, where D614 and G614 variants are visible, (B) hydrophilicity and hydrophobicity analysis of amino acid 611-620, where D614D is shown, (C) hydrophilicity and hydrophobicity analysis of amino acid 611-620, where D614G is shown (D) target sequence 3D model, (E) 3D model of amino acid 589-639; white arrow indicates D614D, (F) 3D model of amino acid 589-639; white arrow indicates D614G.

**Supplementary figure 2:**
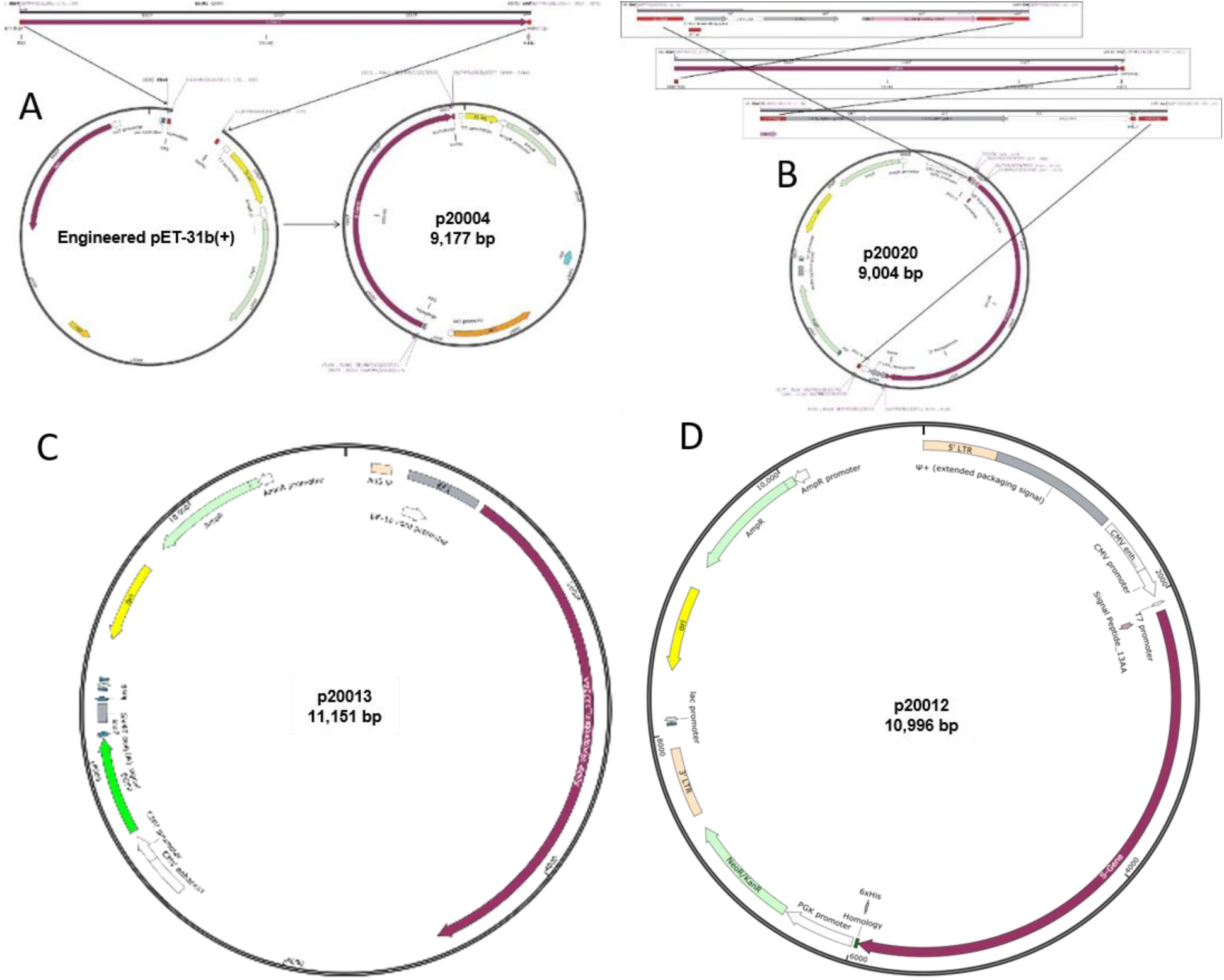
Vector construction and modification. (A) Graphical representation of S-gene and engineered pET31b vector assembly, (B) Graphical representation of p20010 rDNA molecular cloning, (C) Pseudotyped adenoviral rDNA p20013 map, containing EGFP along with S-gene, (D) Pseudotyped retroviral rDNA p20012 map, containing S-gene.

**Supplementary figure 3:**
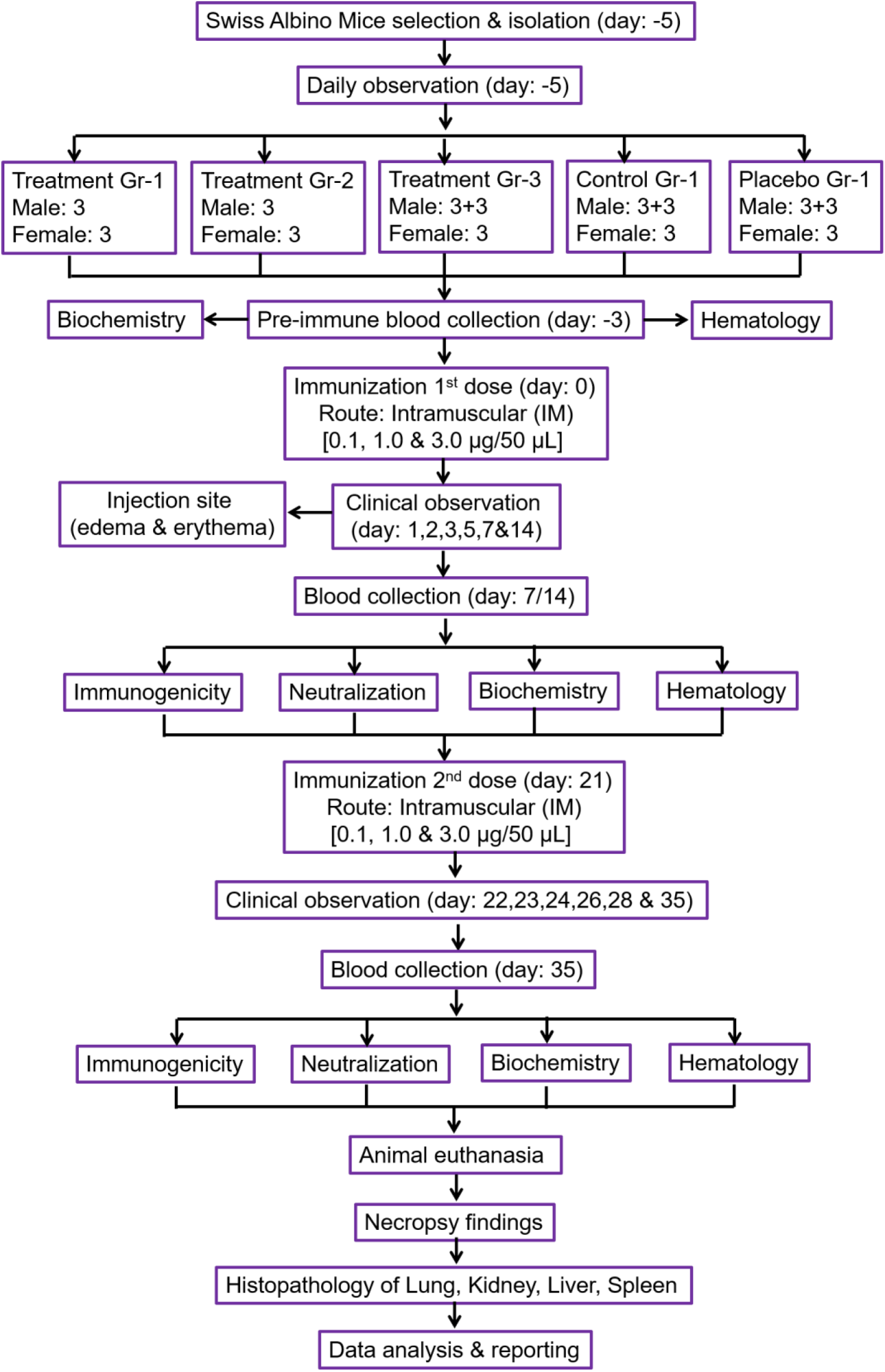
Experimental design for safety and efficacy analysis in mice.

**Supplementary figure 4:**
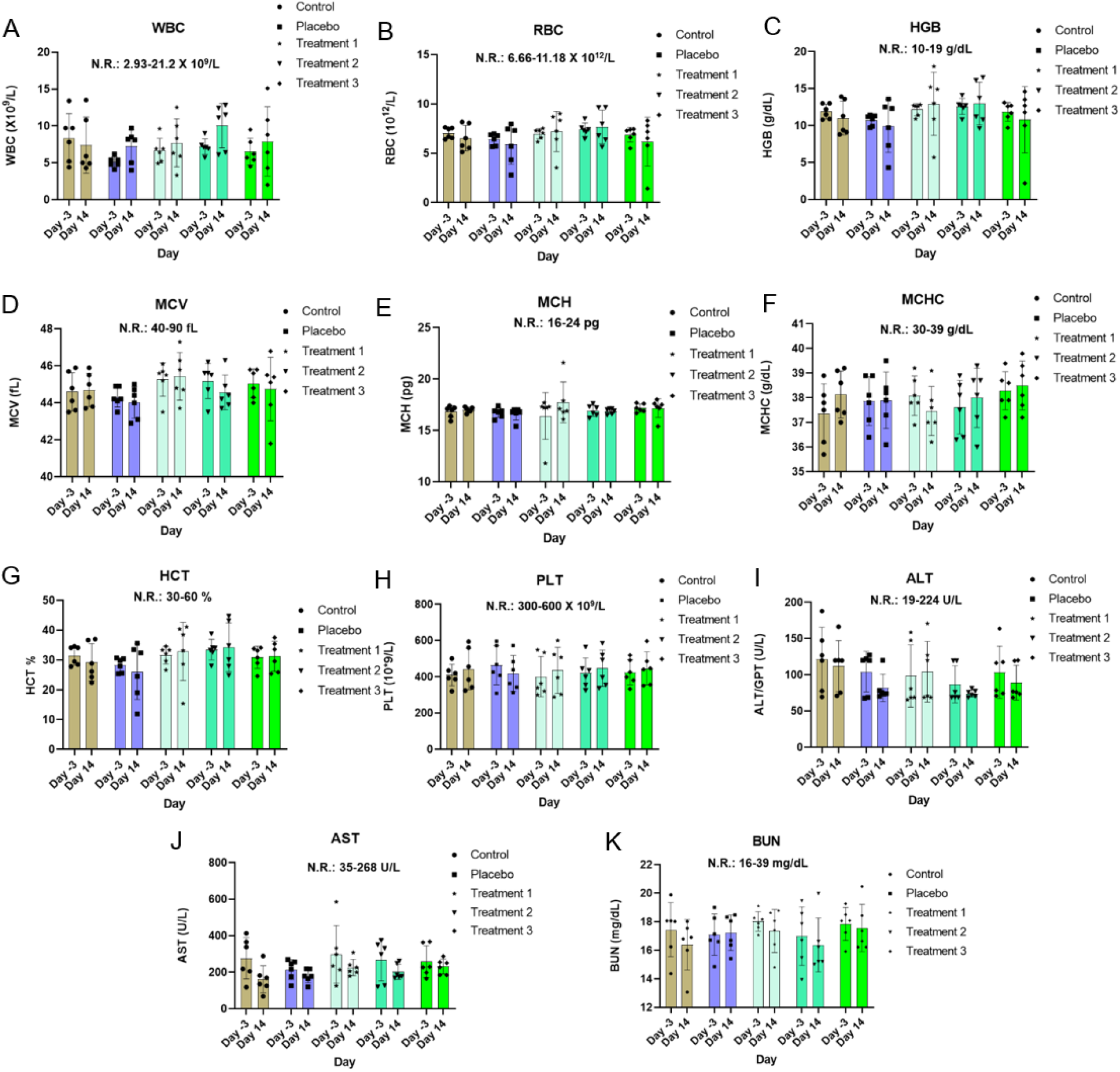
Toxicity analysis, Complete blood count and Chemistry analysis (A) WBC, white blood count (B) RBC, red blood cell (C) MCV, mean corpuscular volume (D) MCHC, mean corpuscular hemoglobin concentration (E) HGB, hemoglobin (F) HCT, hematocrit (G) MCH, mean corpuscular hemoglobin (H) PLT, platelet (I) ALT/GPT, alanine transaminase (J) AST/GOT, aspartate aminotransferase (K) BUN, blood urea nitrogen.

**Supplementary figure 5:**
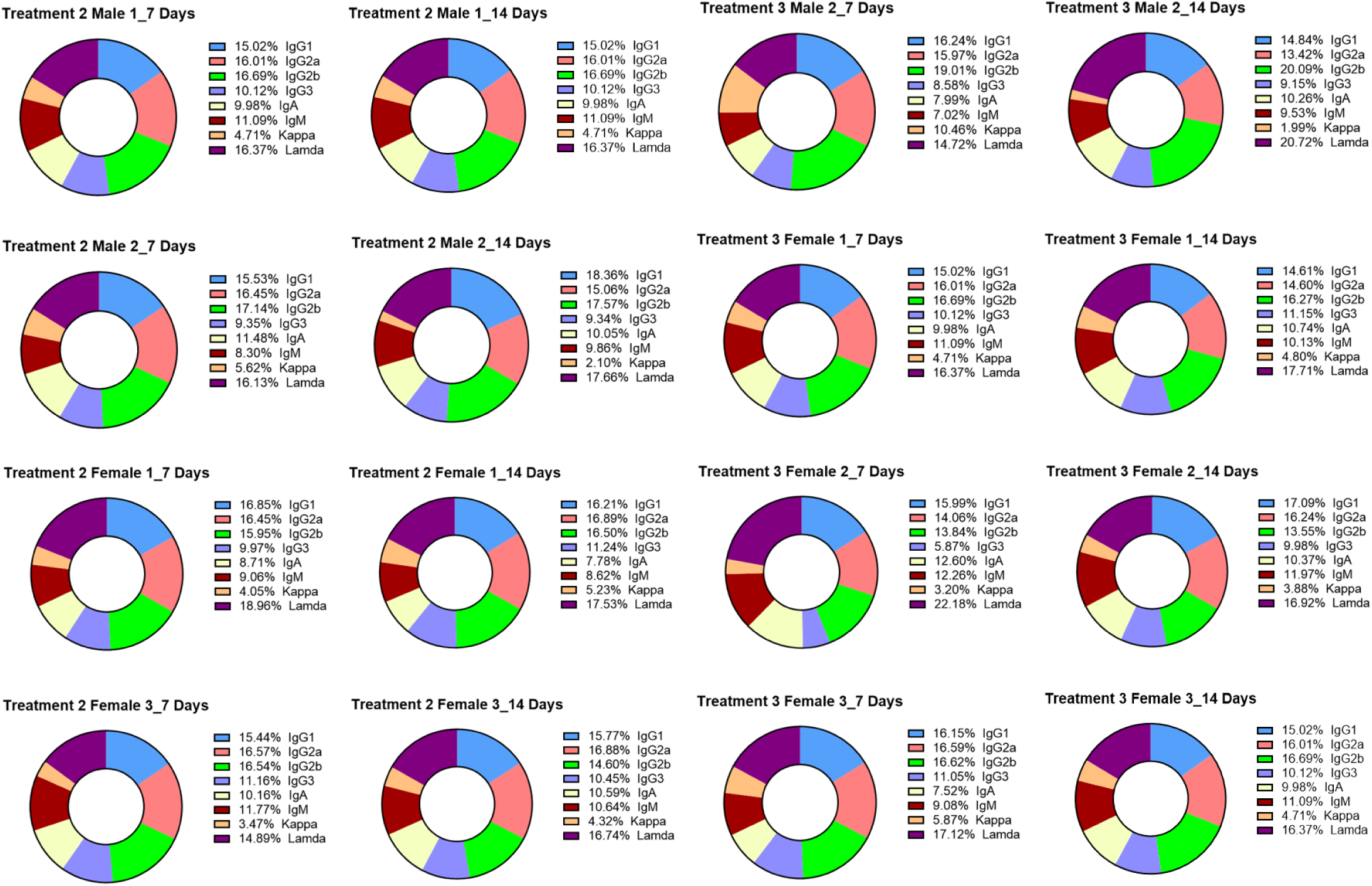
mAb isotyping of representative treatment group of mice, representative sample descriptions are mentioned in respective pie charts.

**Supplementary figure 6:**
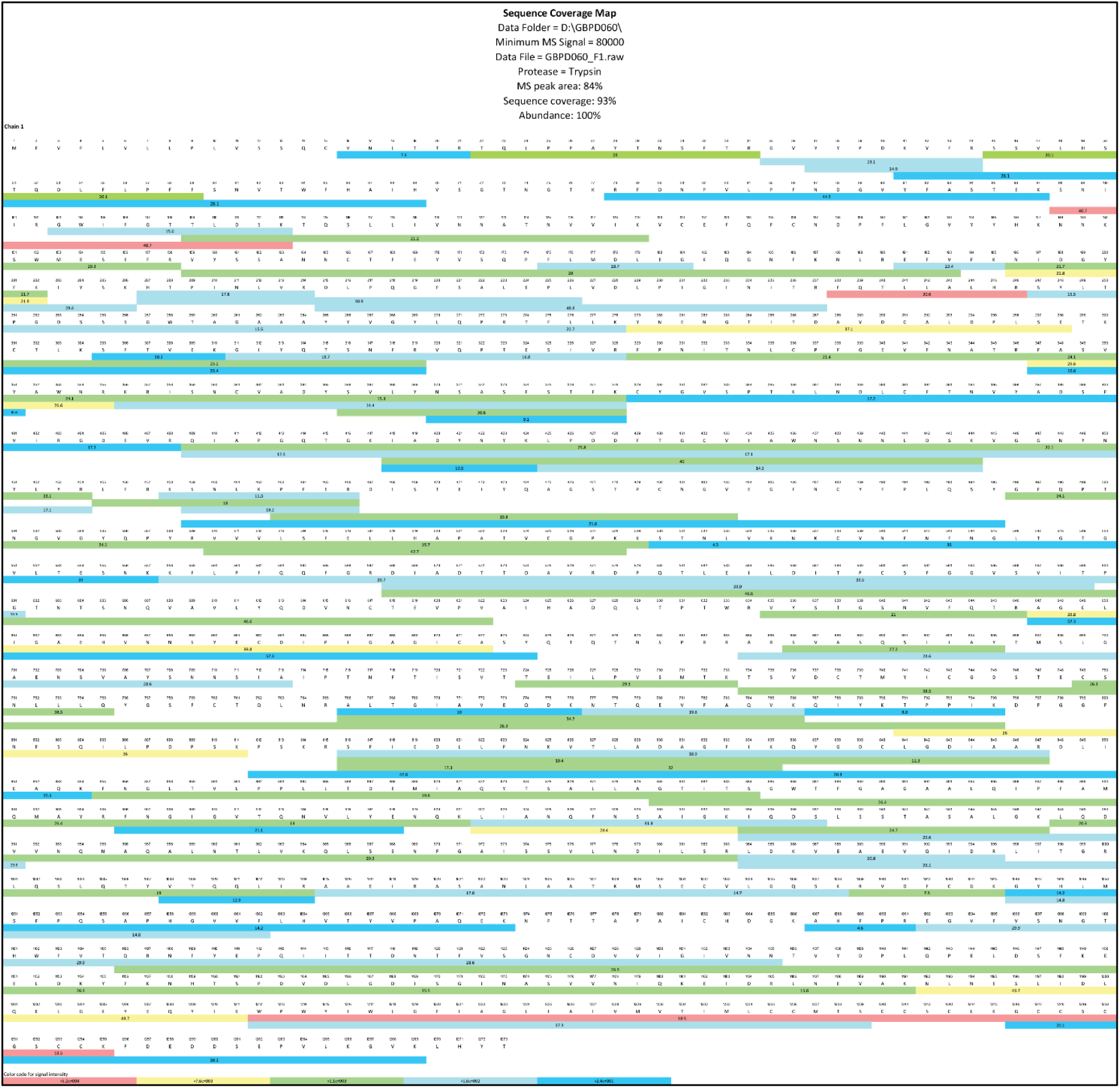
SARS-CoV-2 S protein mapping via LC-MS/MS.

**Supplementary figure 7:**
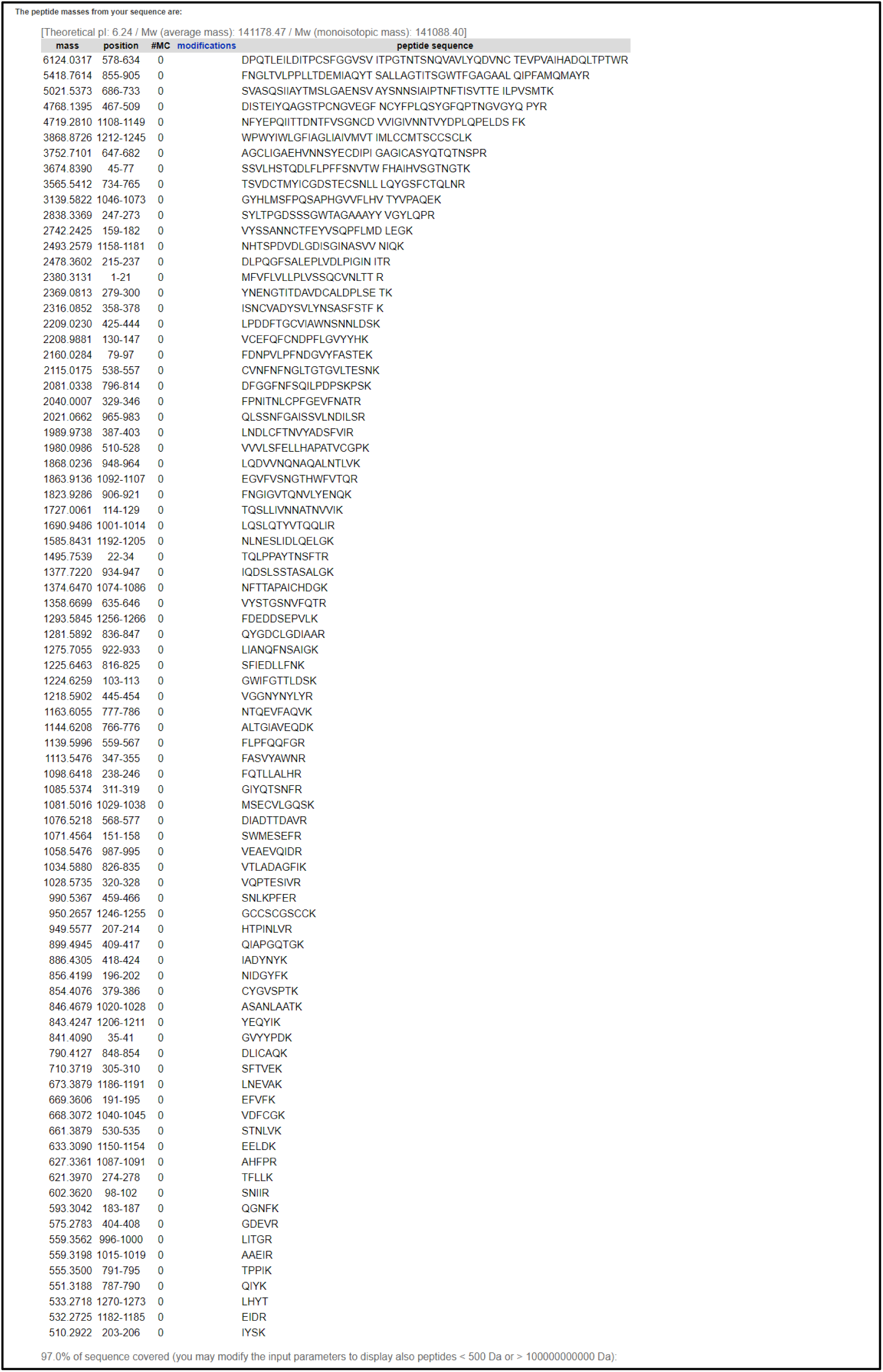
SARS-CoV-2 S protein mapping via ExPASy PeptideMass.

**Supplementary figure 8:**
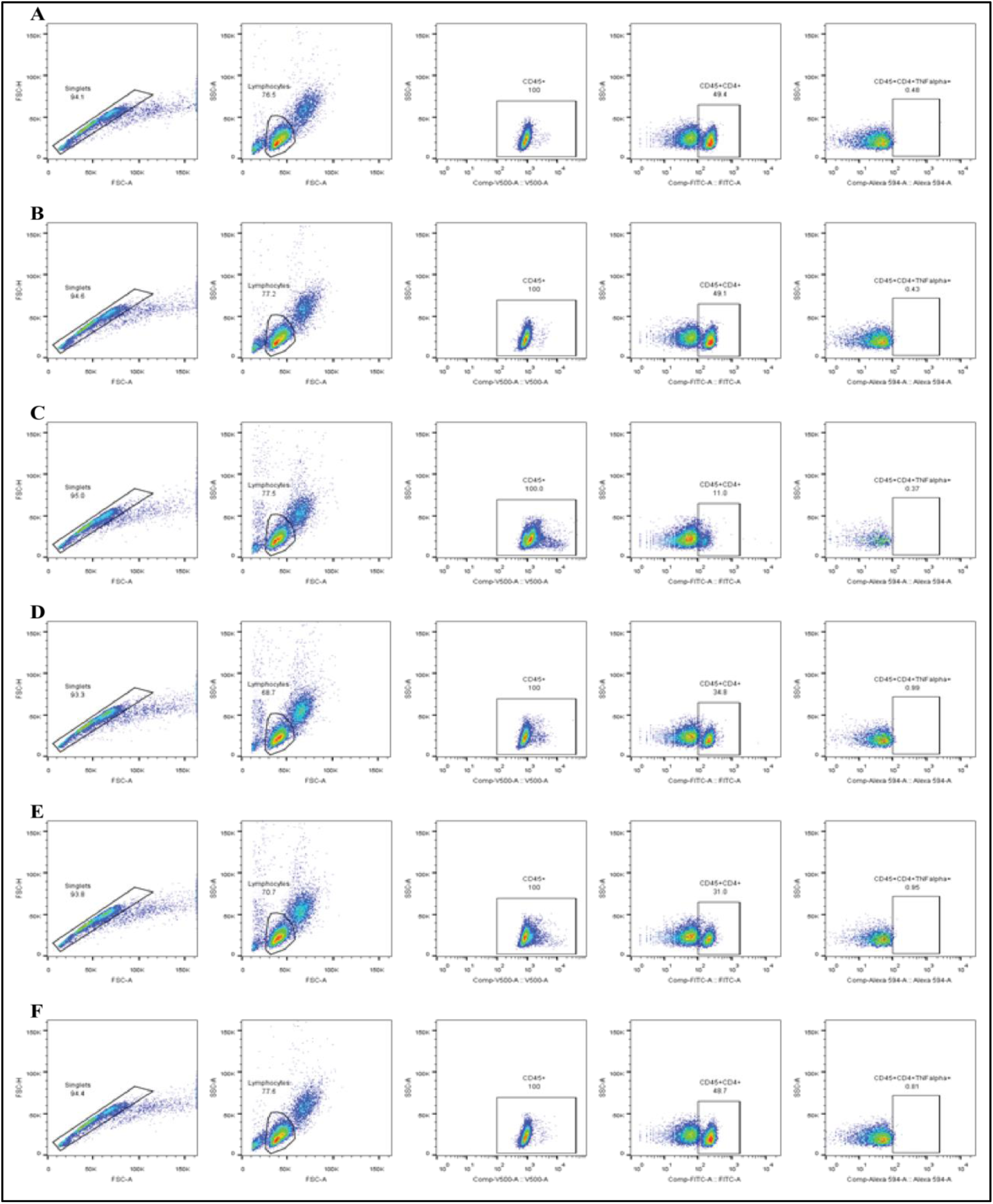
Flow cytometric analysis of total T cell (CD4^+^) populations producing TFN alpha on mouse splenocyte upon SARS-CoV-2 S protein stimulation. Cells were gated in an orderly manner, like singlets were gated, followed by lymphocytes, CD45^+^, CD45^+^CD4^+^ and CD45^+^CD4^+^TFNalpha^+^ (A, B, C) 3 control panels where 0.48%, 0.43% and 0.37% CD45^+^CD4^+^TFNalpha^+^ cells were identified respectively, (D, E, F) 3 treatment panels where 0.99%, 0.95% and 0.81% CD45^+^CD4^+^TFNalpha^+^ cells were identified respectively.

**Supplementary figure 9:**
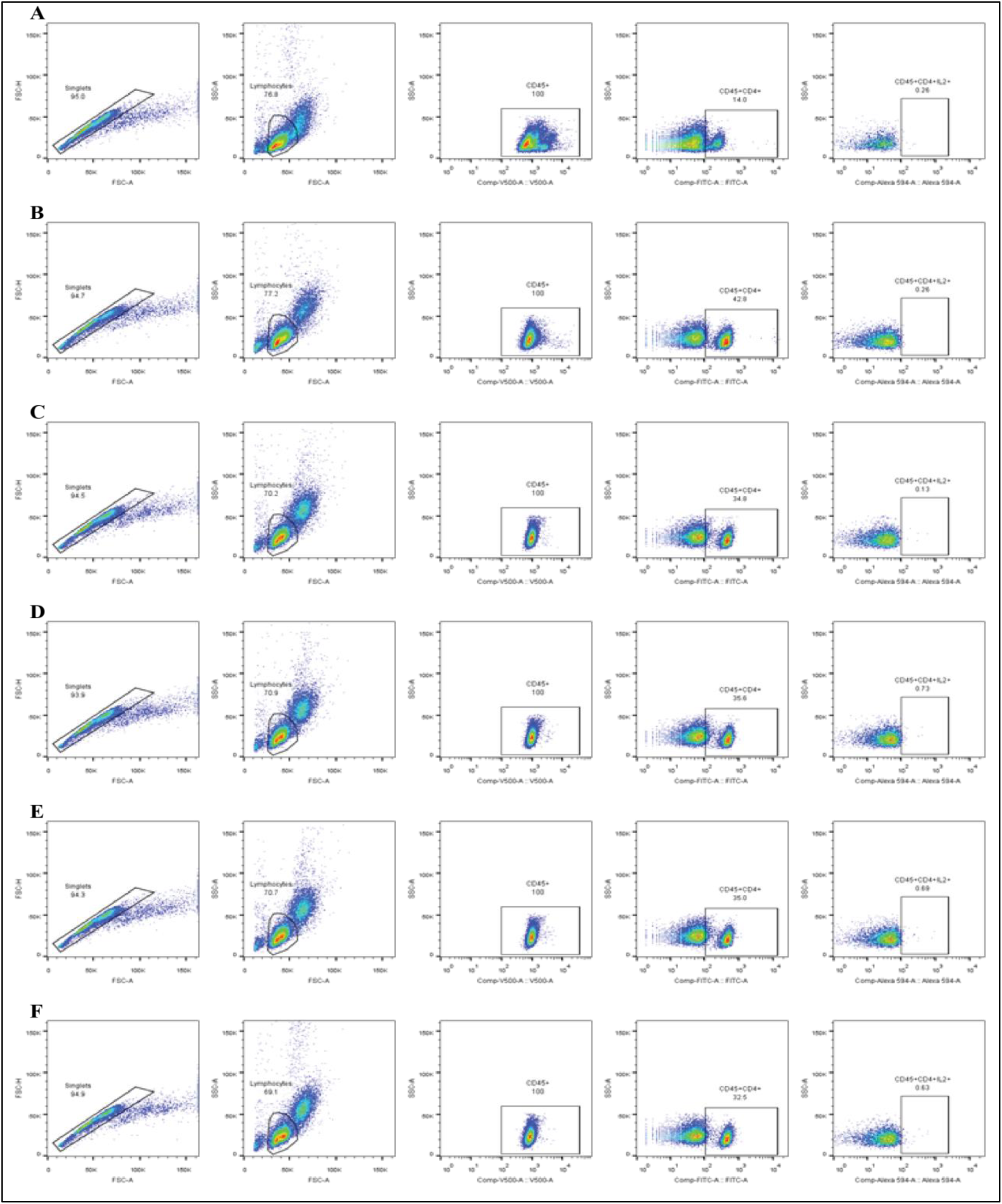
Flow cytometric analysis of total T cell (CD4^+^) populations producing IL-2 on mouse splenocyte upon SARS-CoV-2 S protein stimulation. Cells were gated in an orderly manner, like singlets were gated, followed by lymphocytes, CD45^+^, CD45^+^CD4^+^ and CD45^+^CD4^+^IL2^+^ (A, B, C) 3 control panels where 0.26%, 0.26% and 0.13% CD45^+^CD4^+^IL2^+^ cells were identified respectively, (D, E, F) 3 treatment panels where 0.73%, 0.69% and 0.63% CD45^+^CD4^+^IL2^+^ cells were identified respectively.

**Supplementary figure 10:**
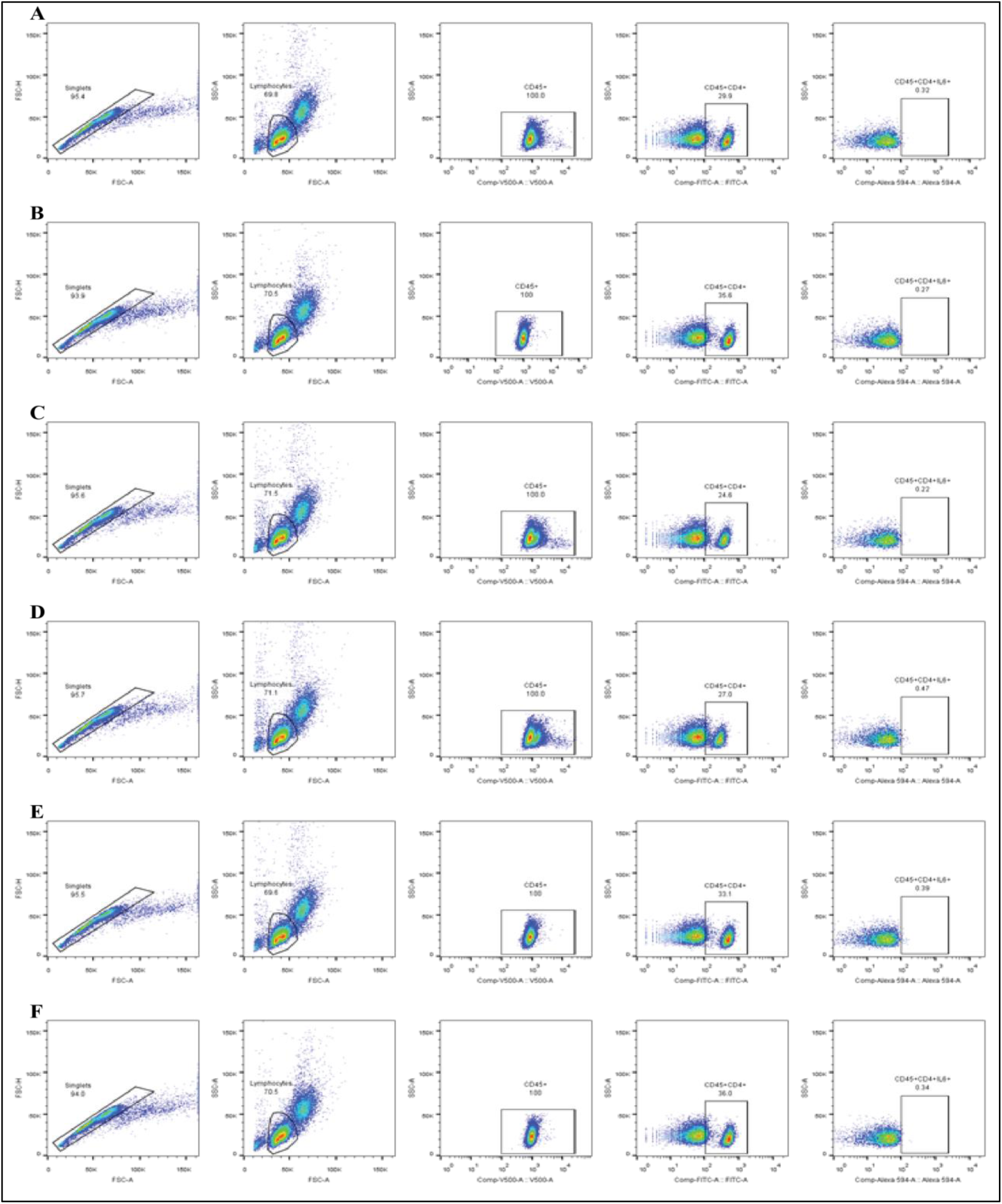
Flow cytometric analysis of total T cell (CD4^+^) populations producing IL-6 on mouse splenocyte upon SARS-CoV-2 S protein stimulation. Cells were gated in an orderly manner, like singlets were gated, followed by lymphocytes, CD45^+^, CD45^+^CD4^+^ and CD45^+^CD4^+^IL6^+^ (A, B, C) 3 control panels where 0.32%, 0.27% and 0.22% CD45^+^CD4^+^IL6^+^ cells were identified respectively, (D, E, F) 3 treatment panels where 0.47%, 0.39% and 0.34% CD45^+^CD4^+^IL6^+^ cells were identified respectively.

**Supplementary table 1:**
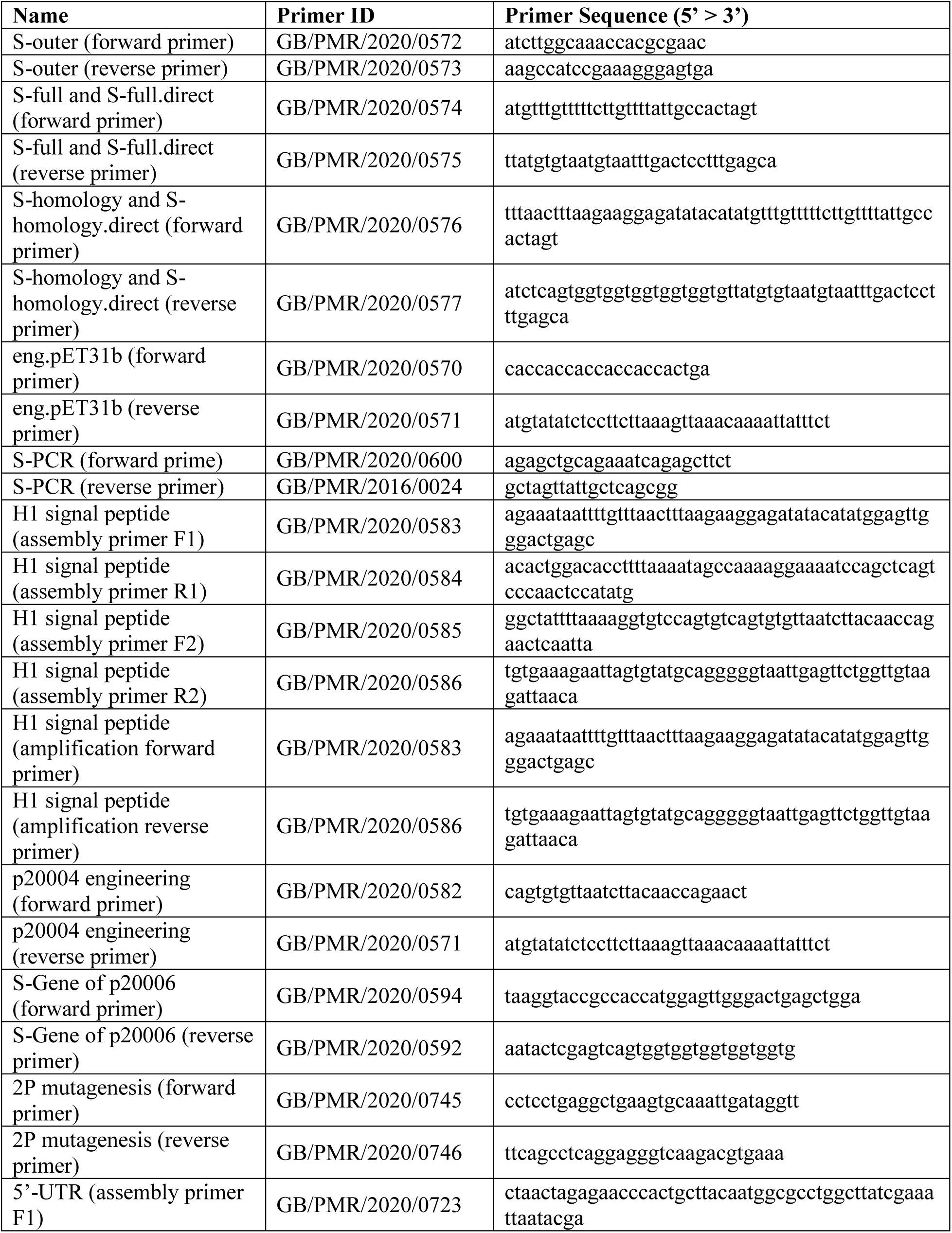
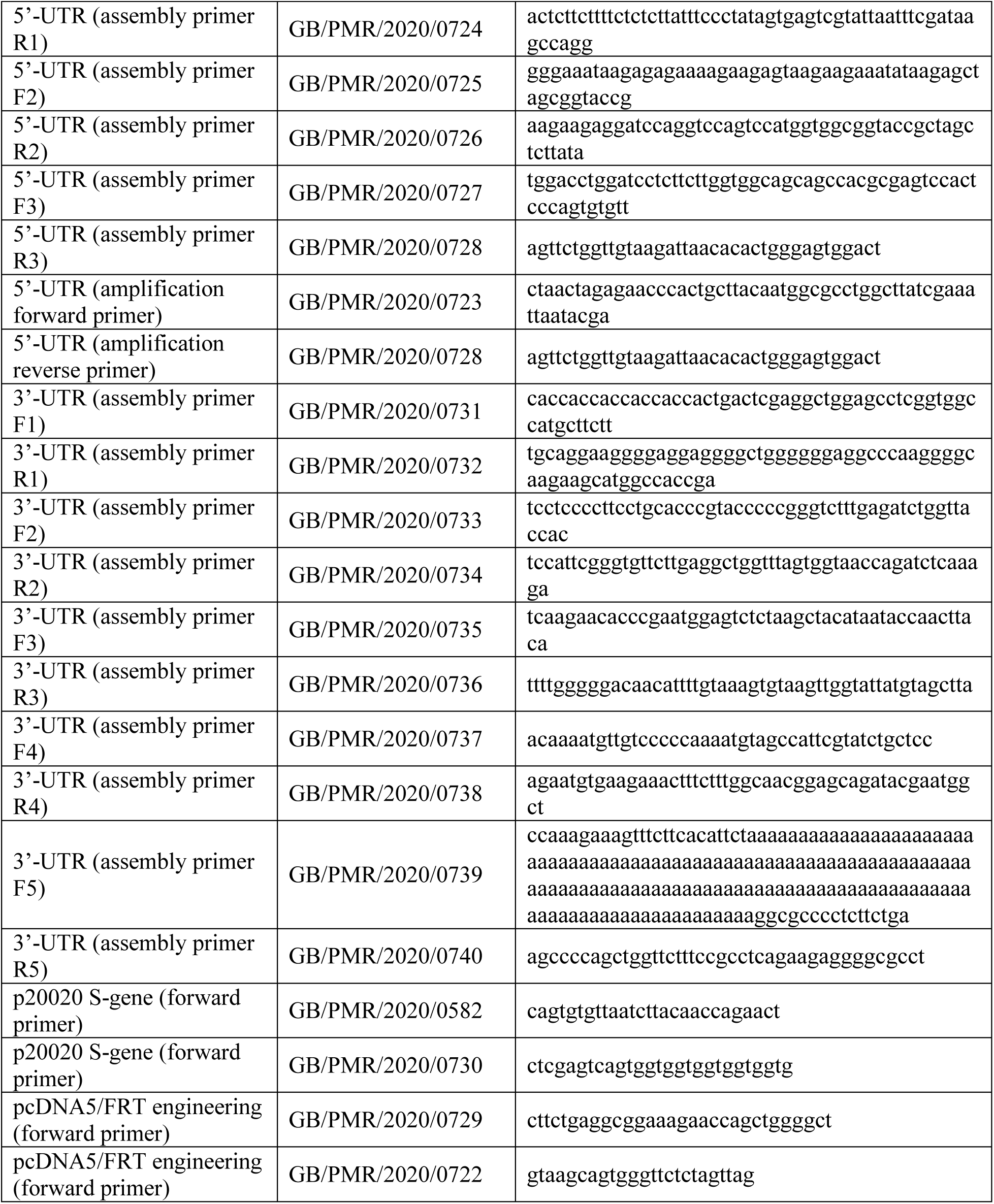
assembly, amplification, engineering and mutagenesis primers

**Supplementary table 2:**
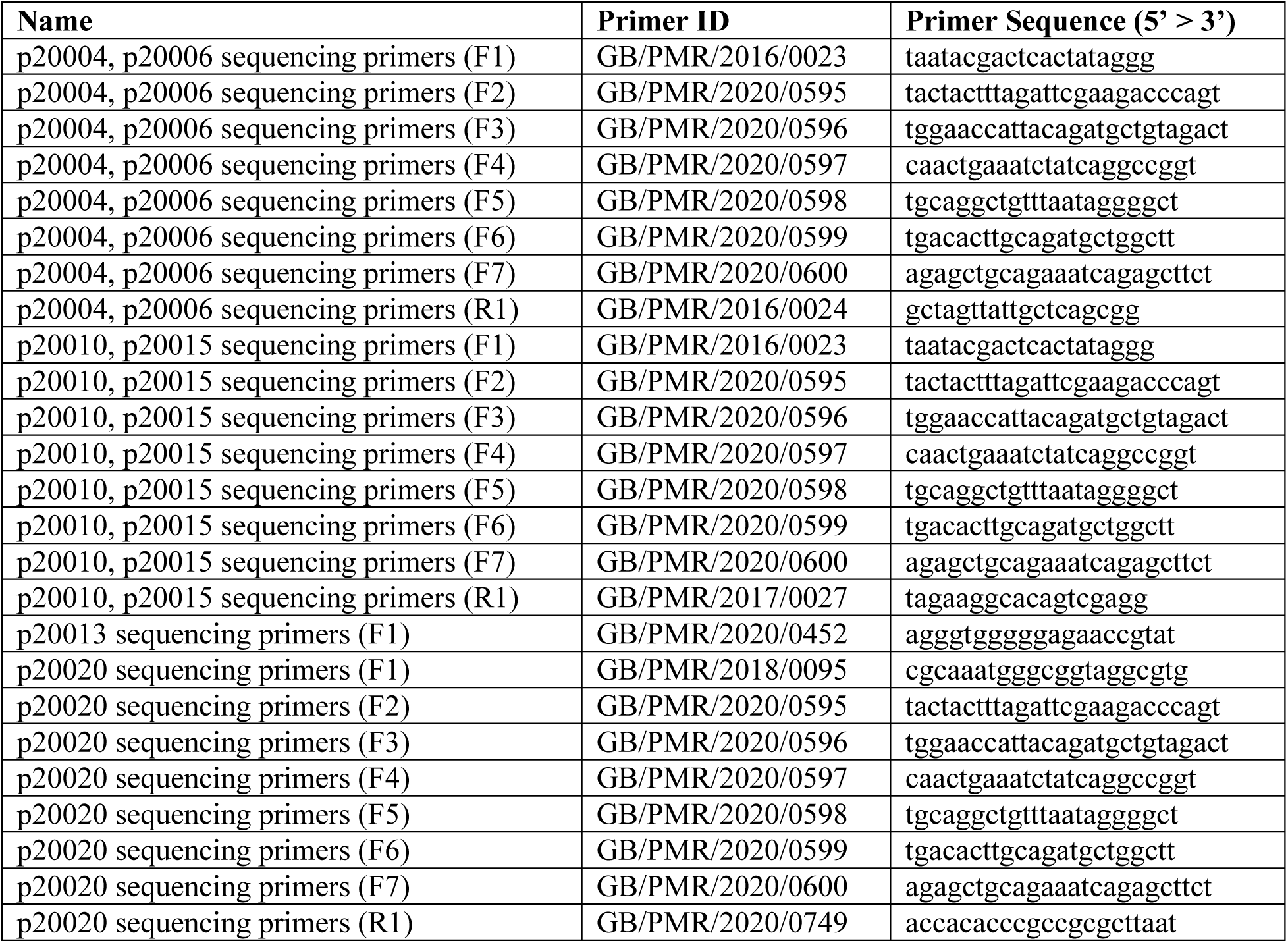
sequencing primers

**Supplementary table 3:**
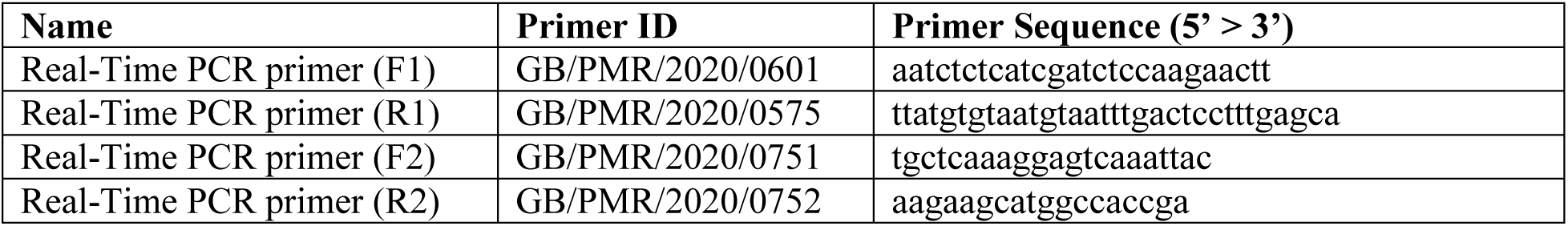
real-time PCR primers

## Supplementary method 1: IVT mRNA synthesis optimization

### DNA preparation for IVT reaction

30 µg of p20020 rDNA was restriction digested with SfoI (ThermoFisher, USA) for 16 hours, visualized using 0.8% agarose gel electrophoresis, gel excised and DNA extracted from gel using GeneJET Gel Extraction and DNA Cleanup Micro Kit, re-purification of DNA by phenol:chloroform:isoamyl alcohol, followed by phenol removal using chloroform (twice). Purified lyophilized DNA was reconstituted using nuclease-free water, quantified and store at -30 °C for future use.

Optimization step 1: Synthesis time factor

240 ng linear purified DNA was used for all 4 optimization reactions. Each reaction was performed in a 20 µL total volume. For every reaction, a DNase treatment reaction was also performed using 1 µL TURBO DNase (2 U/µL) at 37 °C for 15 minutes. For visualization, 1% agarose gel electrophoresis was performed after every step of reaction (**figure 1C**).

In optimization step 1, where synthesis time dependency was observed, for that following components were mixed together apart from water and template, and reaction were run for 2, 4, 6, 8, 10 and 16 hours. 3 control reactions were also performed (1 µg control template pTRI-Xef for each reaction) for 2, 4 and 16 hours at 37 °C.

| Items | Final conc. |
| --- | --- |
| ATP | 7.5 mM |
| UTP | 7.5 mM |
| GTP | 7.5 mM |
| CTP | 7.5 mM |
| 10X reaction buffer | 1X |
| 10X enzyme mix | 1X |

Optimization step 2: rNTPs concentration

In 2^nd^ step of optimization, where rNTPs concentration was observed at a constant synthesis time (2 hours) at 37 °C. For that following components were mixed together apart from water and template. As this point, last optimized condition was run as positive control.

| Items | Final conc. |  |  |  |
| --- | --- | --- | --- | --- |
| ATP | 7.5 mM | 13 mM | 16.5 mM | 21.5 mM |
| UTP | 7.5 mM | 13 mM | 16.5 mM | 21.5 mM |
| GTP | 7.5 mM | 8.0 mM | 10 mM | 13.5 mM |
| CTP | 7.5 mM | 7.5 mM | 9.5 mM | 12.5 mM |
| 10X reaction buffer | 1X | 1X | 1X | 1X |
| 10X enzyme mix | 1X | 1X | 1X | 1X |

Optimization step 3: RNase inhibitor and pyrophosphatase effect

In 3^rd^ step of optimization, murine RNase inhibitor and yeast pyrophosphatase effects were observed at a constant synthesis time (2 hours) and constant rNTPs at 37 °C. For that following components were mixed together apart from water and template. A higher concentration of rNTPs reaction was setup. As this point, last optimized condition was run as positive control.

Supplementary Documents
| Items | Final conc. |  |
| --- | --- | --- |
| ATP | 7.5 mM | 13.13 mM |
| UTP | 7.5 mM | 13.13 mM |
| GTP | 7.5 mM | 9.38 mM |
| CTP | 7.5 mM | 9.38 mM |
| 10X reaction buffer | 1X | 1X |
| 10X enzyme mix | 1X | 1X |
| RNase inhibitor, Murine | 1 U/μL | 1 U/μL |
| Pyrophosphatase, inorganic yeast | 0.002 U/μL | 0.002 U/μL |
| Figure 1C, lane number → | 1 & 2 | 3 & 4 |

Optimization step 4: Temperature dependency

In 4^th^ step of optimization, temperature dependency was observed at a constant synthesis time (2 hours), constant rNTPs, and constant RNase inhibitor and pyrophosphatase, and at 38, 37, 36, 35, 34 and 33 °C. For that following components were mixed together apart from water and template. A higher concentration of rNTPs reaction was also setup. As this point, last optimized condition was run as positive control.

| Items | Final conc. |  |
| --- | --- | --- |
| ATP | 13.13 mM | 21.5 mM |
| UTP | 13.13 mM | 21.5 mM |
| GTP | 9.38 mM | 13.5 mM |
| CTP | 9.38 mM | 12.5 mM |
| 10X reaction buffer | 1X | 1X |
| 10X enzyme mix | 1X | 1X |
| RNase inhibitor, Murine | 1 U/μL | 1 U/μL |
| Pyrophosphatase, inorganic yeast | 0.002 U/μL | 0.002 U/μL |

## Supplementary method 2: peptide pool preparation and purification

### Peptide pool preparation

Dissolve 40 µg of SARS-CoV-2 Spike S1+S2 ECD His recombinant protein (Sino Biological, China), S2 ECD-His Recombinant Protein (Sino Biological, China), and RBD-His Recombinant Protein (Sino Biological, China) in 50mM ammonium bicarbonate (Wako Pure Chemicals Industries Ltd., Japan), pH 8 containing 8M urea (ThermoFisher Scientific, USA). After dissolving, add 500 mM DTT (ThermoFisher Scientific, USA) to the solution to a final concentration of 20 mM (1:25 dilution) and mix briefly; incubate at 60 °C for 1 hour. For alkylation, add 1M IAA (Sigma-Aldrich, USA) solution to the reduced protein sample to a final concentration of 40 mM (1:25 dilution); incubate the reaction mixture for 30 minutes protected from light. To stop the reaction, add 500 mM DTT solution to a final concentration of 10 mM (1:50 dilution). To digest, add trypsin (ThermoFisher Scientific, USA) solution to the sample solution to a final trypsin to protein ratio of 1:23 (w/w). Incubate the sample tube at 37 °C for 16 – 24 hours. After incubation, to stop digestion, add formic acid to lower pH 2.0.

### Peptide pool purification

Tapping C18 spin column (ThermoFisher Scientific, USA) to settle resin. Place column into a receiver tube. To activate the column, add 200 µL 50% acetonitrile (Wako Pure Chemicals Industries Ltd., Japan) to wet resin. Centrifuge the column at 1500 × g for 1 minute. Repeat the step. To equilibrate, add 200 µL 0.5% formic acid (Wako Pure Chemicals Industries Ltd., Japan) in 5% acetonitrile (Wako Pure Chemicals Industries Ltd., Japan). Centrifuge the column at 1500 × g for 1 minute. Repeat the step. Load sample on top of resin bed. Place column into a receiver tube. Centrifuge the column at 1500 × g for 1 minute. To ensure complete binding, recover flow-through and repeat the step 2 – 3 times. To wash the column, Place column into a receiver tube. Add 200 µL 0.5% formic acid in 5% acetonitrile to column. Centrifuge the column at 1500 × g for 1 minute. Repeat the step. To recover sample, place column in a new receiver tube. Add 20 µL 70% acetonitrile to top of the resin bed. Centrifuge at 1500 × g for 1 minute. Repeat the step in same receiver tube.

## Supplementary method 3: Mouse splenocyte isolation

RPMI complete media (RPMI + L-glutamine + penicillin streptomycin + mouse sera) was prepared first. Then a 100 mm petri dish was taken, 10 mL complete media was added and harvested spleen was taken into the dish. By using microscopic glass slides, spleen was smashed into pieces within the petri dish. Cells were washed out from slides using micropipette. A 10 ml pipette was used to draw the solution up and down, each time closing the end of the pipette against the bottom of the petri dish – to forcefully expel the contents and break up the pieces. Cell solution was passed through a sterile 40 μm mesh strainer. Centrifugation was performed for 10 minutes at 250 xg, at 4 °C. Supernatant discarded and cells were re-suspended in RBC(1X) lysing buffer (10X RBC lysis buffer: NH4Cl - 4.01 gm, NaHCO3 - 0.42 gm, EDTA - 0.19 gm, pH adjusted to 7.4 using NaOH, volume adjusted to 50 ml with water. Filter sterilize and store at 4 °C for six months.) and incubated at room temp for 3-5 mins. Vigorous shaking was performed at 1 minute intervals. Again centrifugation was performed for 10 minutes at 250 xg, at 4 °C. Supernatant discarded and cells were resuspended in PBS, following centrifugation and supernatant discard. PBS washing step was repeated again. Finally, re-suspension of cell pellet in 3 ml RPMI complete media, plating in a 6-well culture plate and incubate at 37°C, 5% CO2 as needed.

## Notes

### Competing Interest Statement

The authors have declared no competing interest.

